# Dual-site specificity of the archaeal tRNA m^2^G methyltransferase Trm14

**DOI:** 10.64898/2026.08.09.743744

**Authors:** Teppei Matsuda, Takashi Yokogawa, Soichiro Hidetaka, Manaka Sora, Aoi Ihara, Ayumu Toba, Kumpei Kawai, Go Norimoto, Akira Hirata, Hiroyuki Hori, Ryota Yamagami

**Affiliations:** Department of Applied Chemistry, Graduate School of Science and Engineering, Ehime University, Japan; Department of Chemistry and Biomolecular Science, Faculty of Engineering, Gifu University, Japan; Center for One Medicine Innovative Translational Research (COMIT), Gifu University, Japan; United Graduate School of Drug Discovery and Medical Information Sciences, Gifu University, Japan; Department of Natural Science, Division of Science and Technology, Graduate School of Sciences and Technology for Innovation, Tokushima University, Japan

**Keywords:** Archaea, tRNA, tRNA methyltransferase, tRNA methylation, dual-site specificity

## Abstract

*N*^2^-methylguanosine (m^2^G) is widely found at multiple positions in tRNAs across the three domains of life. Tryptophan tRNA from *Thermococcus kodakarensis* contains m^2^G at position 67. We previously proposed that the tRNA m^2^G methyltransferase Trm14 is responsible for m^2^G67 formation in tRNA^Trp^ from *T. kodakarensis*, although Trm14 was originally identified as the enzyme catalyzing m^2^G6 formation in tRNA^Cys^ in *Methanocaldococcus jannaschii*. Thus, it remained unclear whether Trm14 could also methylate G67. Here, we characterized archaeal Trm14. Biochemical analyses using recombinant *T. kodakarensis* Trm14 revealed that the enzyme catalyzes m^2^G formation at positions 6 and 67 in *T. kodakarensis* tRNA^Cys^ and tRNA^Trp^ transcripts, respectively. Mass spectrometric analyses demonstrated the loss of m^2^G6 and m^2^G67 in native tRNA^Cys^ and tRNA^Trp^, respectively, from a *T. kodakarensis trm14* gene disruptant strain, providing direct evidence for the dual-site specificity of *T. kodakarensis* Trm14. The growth phenotype of the *trm14* gene disruptant strain was comparable to that of the wild-type strain. In contrast, a *trm14*/*trm11* double disruptant, in which *trm11* encodes the tRNA m^2^G10/m^2^_2_G10 methyltransferase, exhibited severe growth retardation at 95 °C. This suggests that m^2^G6/m^2^G67 and m^2^G10/m^2^_2_G10 cooperatively contribute to cellular fitness at high temperatures. Biochemical analyses revealed that Trm14 methylates all 46 *T. kodakarensis* tRNA transcripts. Furthermore, we found that recombinant *M. jannaschii* Trm14 methylated both positions. In contrast, the bacterial ortholog TrmN modified only position 6 in tRNA. Overall, this study expands our understanding of archaeal Trm14 by demonstrating its broader substrate specificity and the physiological significance of these modifications under hyperthermophilic conditions.

## Introduction

Transfer RNAs (tRNAs) play essential roles in decoding the genetic information encoded in messenger RNAs during protein synthesis. Beyond translation, tRNAs participate in diverse biological processes, including cell proliferation and human diseases (1–3). Transfer RNAs undergo extensive post-transcriptional modifications that are essential for their proper function. To date, more than 100 distinct modified nucleosides have been identified in tRNAs from various organisms (4). In particular, tRNAs from hyperthermophiles contain numerous modified nucleosides at multiple positions (5,6). While tRNA modifications regulate decoding functions and stabilize tRNA structure (7,8), the stabilization effect on the L-shaped tRNA structure is thought to be especially important in hyperthermophiles for maintaining tRNA functionality at high temperatures (9). However, three important questions remain: which enzymes are responsible for these modifications, how they determine modification sites, and how tRNA modifications contribute to cellular fitness. To address these questions, we focused on *N*^2^-methylguanosine (m^2^G) (Fig. 1A) and *N*^2^, *N*^2^-dimethylguanosine (m^2^_2_G) (Fig. 1B) in archaeal tRNAs in this study.

**Figure 1.**
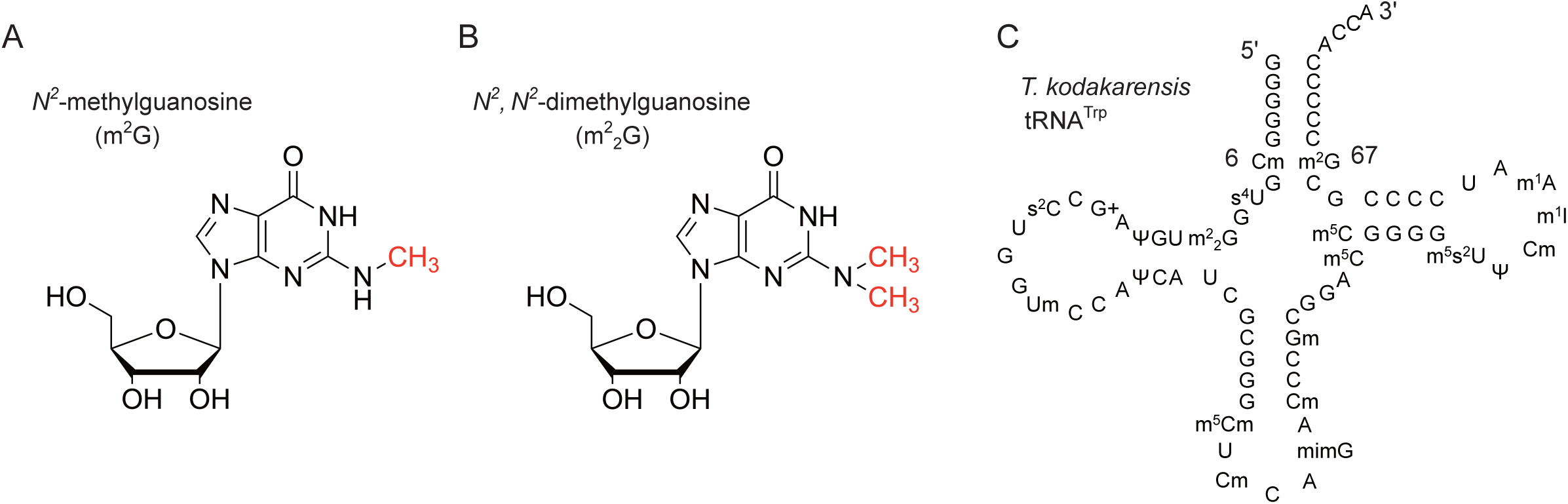
Chemical structures of m^2^G and m^2^_2_G and the cloverleaf structure of *T. kodakarensis* tRNA^Trp^. (A and B) Chemical structures of (A) m^2^G and (B) m^2^_2_G are depicted. Methyl groups are highlighted in red. (C) Primary sequence of *T. kodakarensis* tRNA^Trp^ is shown in the canonical cloverleaf structure.

The m^2^G/m^2^_2_G modifications are ubiquitous modifications that have been identified at positions 6, 7, 9, 10, 18, 26, 27, 57, and 67 in tRNAs from eukaryotes, archaea and eubacteria (10,11). These modifications are installed by site-specific tRNA (m^2^G/m^2^_2_G) methyltransferases. In the case of m^2^G at position 6, the tRNA (m^2^G6) methyltransferase (Trm14) catalyzes the reaction in archaea (12). Trm14 was first identified in *Methanocaldococcus jannaschii* through a bioinformatics-based approach. While the gene was originally annotated as an *N*^6^-adenine-specific DNA methylase, it possesses a THUMP domain, an RNA-binding domain, commonly found in RNA methyltransferases, tRNA thiouridine synthetases, and pseudouridine synthases (13,14). Biochemical analyses using recombinant Trm14 demonstrated that Trm14 catalyzes m^2^G6 formation in tRNA^Cys^ in *M. jannaschii* (12). In bacteria, TrmN, a bacterial ortholog of Trm14, is responsible for m^2^G6 formation in *Thermus thermophilus* tRNA^Phe^ (15,16). The Trm14 and TrmN proteins consist of the following domains: an N-terminal ferredoxin-like domain, THUMP domain, and a C-terminal Rossmann-fold methyltransferase (RFM) domain, which utilizes S-adenosyl-L-methionine (SAM) as a methyl group donor. Furthermore, X-ray crystal structures of these methyltransferases revealed that the overall structures of these enzymes are highly similar, with a root-mean-square deviation (RMSD) of 2.0 Å over 250 Cα atoms. Local structural differences, however, are observed in THUMP domain, where additional antiparallel β-strands are inserted in Trm14 (17). Moreover, the tRNA-binding surface of Trm14/TrmN is formed by both the THUMP and RFM domains. Based on these observations, a tRNA-binding mode of TrmN was proposed (17). A later study identified human THUMPD3-TRMT112 as an enzyme complex responsible for the m^2^G modification at position 6 as well as at position 7 in tRNA of higher eukaryotes (18). Unlike Trm14 or TrmN, THUMPD3 alone is catalytically inactive and requires a small subunit protein, TRMT112 (also known as Trm112 in yeast), to promote SAM binding and tRNA binding, and to stabilize the THUMPD3 structure (18). In addition, Trm112 acts as a hub protein interacting with other methyltransferases and thus plays central roles in regulating gene expression (10,19–23).

The other major m^2^G/m^2^_2_G sites found in tRNA are guanosines at position 10 and 26 (6). In archaea, these sites are methylated by archaeal tRNA (m^2^G10/m^2^_2_G10) methyltransferase (Trm11) as well as tRNA (m^2^G26/m^2^_2_G26) methyltransferase (Trm1) (24–30). While the domain architecture of archaeal Trm11 is also composed of THUMP fused with NFLD and RFM, Trm1 has a distinct domain architecture and lacks a THUMP domain (14,29). In the case of *Archaeoglobus fulgidus*, Trm11 interacts with Trm112, which enhances the catalytic activity (31). Importantly, m^2^G can still participate in the formation of a Watson–Crick base pair with cytidine, whereas m^2^_2_G disrupts canonical G-C base pairing. In the structure of *S. cerevisiae* tRNA^Phe^, the m^2^G10 forms a triple base pair with C25 and G45, which further stacks with m^2^_2_G26- A44 tertiary base pair (32). In addition, we recently found that m^2^_2_G26 modification contributes to tRNA folding into the cloverleaf structure (33). Thus, these m^2^G/m^2^_2_G modifications stabilize the tRNA structure.

*Thermococcus kodakarensis* is a hyperthermophilic archaeon that grows at temperatures ranging from 60 °C to 100 °C (34). In 2019, we determined the primary sequence of tRNA^Trp^ from *T. kodakarensis* by liquid chromatography-tandem mass spectrometry (LC–MS/MS) (Fig. 1C) and found that it contains 21 modified nucleosides among 78 total nucleotides, representing an exceptionally high modification density (35). Our gene orthology analysis identified candidate enzymes responsible for these modifications, and some of the corresponding genes were characterized by reverse genetic analyses (35). For example, the m^2^_2_G at position 10 in tRNA^Trp^ was completely lost upon disruption of the gene encoding the tRNA m^2^G10/m^2^_2_G10 methyltransferase (archaeal *trm11*; Tk0981) in *T. kodakarensis* genome, suggesting that m^2^_2_G10 formation is catalyzed by archaeal Trm11 (35). Importantly, deletion of *trm11* caused growth retardation at high temperatures, demonstrating the physiological significance of m^2^G10/m^2^_2_G10 for cellular fitness in hyperthermophiles (35). Similarly, the 4- thiouridine (s^4^U) at position 8, 5-methyl-2-thiouridine (m^5^s^2^U) at position 54, and methylwyosine (mimG) at position 37 also disappeared upon disruption of tRNA s^4^U synthetase gene (*thiI*; Tk0366), a methyltransferase gene (*rumA*; Tk2135), and *tyw1* gene (Tk1671), respectively (35). More recently, we identified that tRNA (Cm6) methyltransferase TrmTS (Tk1257) is responsible for 2’-*O*-methylcytidine at position 6 in tRNA^Trp^ (36). In addition to the m^2^_2_G10 and m^2^_2_G26 modifications, *T. kodakarensis* tRNA^Trp^ contains an m^2^G modification at position 67 (Fig. 1C) (35). While Trm14 catalyzes m^2^G6 formation in *M. jannaschii*, we previously proposed that archaeal Trm14 is responsible for both m^2^G6 and m^2^G67 formation. However, direct biochemical evidence demonstrating that Trm14 catalyzes m^2^G67 formation is still lacking. Furthermore, the physiological roles of m^2^G6 and m^2^G67 in tRNAs in hyperthermophiles remain unknown. In this study, we addressed these questions through comprehensive biochemical and genetic analyses of archaeal Trm14.

## Materials and Methods

### Materials

[Methyl-^3^H]- S-adenosyl-L-methionine (SAM) (2.47 TBq/mmol) and [Methyl-^14^C]- SAM (1.95 GBq/mmol) were purchased from Revvity. DNA oligonucleotides were obtained from Thermo Fisher Scientific. The *T. kodakarensis* strains and plasmids used in this study are listed in Supplementary Table 1. Primers used in this study are listed in Supplementary Table 2. Gelrite was purchased from FUJIMFILEM WAKO Chemicals. Unless otherwise specified, chemicals were obtained from Nacalai Tesque.

### Strains, media, and culture conditions

*Thermococcus kodakarensis* KUW1 (WT) strain (37) and its derivative gene disruptant strains were cultivated anaerobically at 85 - 95 °C in either nutrient-rich MA-YT medium or synthetic ASW-AA medium. MA-YT medium (1 L) contained 0.8 × Marine Art SF1 reagent (Osaka Yakken Co. Ltd.), 5 g yeast extract (Y), and 5 g tryptone (T). Prior to cultivation, either 5 g sodium pyruvate (Pyr) or 2 g elemental sulfur (S^0^) was added per liter of MA-YT medium. ASW-AA medium contained a vitamin mixture, modified Wolfe’s trace mineral solution and the 20 canonical amino acids dissolved in 0.8 × artificial seawater. Elemental sulfur (S^0^; 2 g L^−1^) was added before inoculation (38,39). All liquid media were supplemented with resazurine (0.5 mg L^−1^) as an oxygen indicator, and 5% (w/v) Na_2_S·9H_2_O was added until the medium became colorless. For colony isolation, ASW-AA media was solidified with 1% (w/v) Gelrite and supplemented with 0.4% (w/v) polysulfide per 100 mL.

### Plasmid construction

*Thermococcus kodakarensis trm14* gene (Tk1863) was PCR amplified from genomic DNA with a set of primers (Supplementary Table 2) and cloned into the NdeI and BamHI sites of pET30a using NEBuilder HiFi DNA Assembly Master Mix (New England Biolabs). The sequence of *M. jannaschii trm14* gene (MJ0438) was designed with codons optimized for expression in *E. coli* system and synthesized by GenScript (Supplementary Table 2). The synthesized gene was cloned into the NcoI and BamHI sites of pE-SUMOpro Amp vector (LifeSensors), using NEBuilder HiFi DNA Assembly Master Mix. The plasmid encoding the *trmN* gene from *Thermus thermophilus* HB27 was constructed previously (16).

### Expression and purification of recombinant proteins

*E. coli* BL21 (DE3) Rosetta 2 cells transformed with the expression plasmids were cultured in 100 mL of LB medium supplemented with 50 µg/ml kanamycin and 30 µg/ml chloramphenicol for *T. kodakarensis* Trm14 or 100 µg/ml ampicillin and 30 µg/ml chloramphenicol for *M. jannaschii* Trm14 at 37 °C for 12-16 hours. The cultured cells were then inoculated into 1 L of LB medium and further grown at 37 °C. When the optical density at 600 nm reached ∼0.8, isopropyl-ß-D-thiogalactopyranoside (IPTG) was added to a final concentration of 1 mM. Protein expression was induced at 37 °C for 4 h. The cells were collected by centrifugation at 10,000 x g at 4 °C for 20 min. One gram of wet-cells expressing *T. kodakarensis* Trm14 was resuspended in buffer A containing 50 mM Tris-HCl (pH 7.6), 1 mM EDTA, 100 mM KCl, and 6 mM 2- mercaptoethanol supplemented with 1x protease inhibitor solution. Cells were disrupted with an ultrasonic disruptor (model VCX-500, Sonics & Materials, Inc.) on ice for 30 min with a pulse interval of 3 sec ON and 2 sec OFF. Cell debris was removed by centrifugation at 10,000 x g at 4 °C for 20 min. The supernatant was incubated at 75 °C for 30 min, and precipitated proteins were removed by centrifugation at 10,000 x g at 4 °C for 20 min. The supernatant was loaded onto a HiTrap Q HP (5 mL; Cytiva) column pre-equilibrated with buffer A. After washing with 25 mL buffer A, Trm14 was eluted with a linear KCl gradient from 200 mM to 1000 mM in 50 mL buffer A. Fractions containing Trm14 were pooled and loaded onto a HiTrap Heparin HP column pre-equilibrated with buffer A. After washing with 25 mL buffer A, Trm14 was eluted with a linear KCl gradient from 200 mM to 1000 mM in 50 mL buffer A. The purified Trm14 was concentrated to 3.5 mg/mL using Vivaspin 15R centrifugal filter units (Sartorius) and stored in a storage buffer containing 50 mM Tris-HCl (pH 7.6), 100 mM KCl, 6 mM 2-mercaptoethanol, and 50% (v/v) glycerol at -80 °C. For purification of *M. jannaschii* Trm14, the heat denaturation step was performed at 70 °C instead of 75 °C, whereas all other purification procedures were identical. *T. thermophilus* TrmN was purified as described previously (16).

### Preparation of tRNA transcripts

RNA transcripts were prepared by in vitro transcription as described previously (40). The oligonucleotides used in this study were provided in Supplementary Table 3.

### In vitro methylation assay of archaeal Trm14 and *T. thermophilus* TrmN

In vitro reaction of Trm14 was performed in a 160 or 70 µL reaction mixture containing 50 mM Tris-HCl (pH 7.6), 50 mM KCl, 5 mM MgCl_2_, 6 mM 2-mercaptoethanol, 20 µM tRNA, 100 nM [Methyl-^3^H]-SAM and 3 µM *T. kodakarensis* or *M. jannaschii* Trm14.

Twenty or ten microliters of the mixture were taken at various time points (for *T. kodakarensis* Trm14; 0, 1, 5, 10, 30, 45, and 60 min, and for *M. jannaschii* Trm14; 0, 2, 5, 10, 30, and 60 min) and spotted onto a Whatman 3MM filter. In the case of *T. thermophilus* TrmN, the reaction was performed at 60 °C. Five microliters of the reaction mixture were taken at various time points (0, 2, 5, 10, 20, 30, and 60 min) and spotted onto a Whatman 3MM filter. The filters were washed with chilled 5% (w/v) trichloroacetic acid eight times at 4 °C for 5 min and then dried. Methyl group incorporation was measured by a liquid scintillation counter.

### Purification of native tRNAs from *T. kodakarensis*

Total RNA was purified as described previously (36). Total RNA was separated by 10% denaturing PAGE (7 M urea), and the tRNA fraction was excised. tRNAs were eluted from the gels in 400 µL of TEN250 buffer containing 10 mM Tris-HCl (pH 7.6), 1 mM EDTA, and 250 mM NaCl, followed by filtration to remove gel debris. The tRNA fraction was recovered by ethanol precipitation. Transfer RNA^Trp^ and tRNA^Cys^ were further isolated from the tRNA fraction using the solid-phase DNA probe method (41,42). The sequences of the DNA probes were provided in Supplementary Table 4.

### LC–MS/MS analysis of tRNA fragments

For full-length RNA, RNA was diluted with ultrapure water to a final concentration of approximately 50 µg/mL as a sample. To obtain RNA fragments, 1 µg of full-length RNA and either 0.5 µg of RNase A or 2 µg of RNase T1 were added to 10 µL of 10 mM triethylamine bicarbonate (TEAB, pH 7.6). The reaction mixture was incubated at 37 °C for 2 h, followed by incubation at 65 °C for 5 min, and then further incubated at 37 °C for an additional 2 h. After the reaction, RNase was inactivated by phenol–chloroform extraction. The RNA fragment was then adsorbed onto 10 µL of Q-Sepharose HP resin, washed five times with 200 µL of 10 mM TEAB (pH 7.6), and eluted with 50 µL of 1 M TEAB (pH 7.6). The eluate was dried using a centrifugal evaporator and subsequently dissolved in 20 µL of ultrapure water.

For analysis, a Shimadzu Nexera XS inert HPLC system was used with water as mobile phase A and 90:10 methanol/water (v/v) as mobile phase B, both with 8 mM triethylamine and 100 mM 1,1,3,3,3-hexafluoroisopropanol. A sample (5–10 µL) was injected into a CAPCELL CORE C18 column (φ2.1 × 100 mm, 2.7 µm particle size; OSAKA SODA). RNA fragments were developed at 60 °C at a flow rate of 0.3 mL/min with a gradient as follows: 100% mobile phase A from 0 to 2 min, 0–100% mobile phase B from 2 to 8 min, 100% mobile phase B from 8 to 10 min and 100% mobile phase A from 10 to 11 min. To analyze mass, a SCIEX ZenoTOF 7600 system was used in negative polarity using a TOF-MS mode for measuring parent mass and MRM^HR^ mode for measuring collision-induced dissociation. The operating parameters were as follows: ion source gas 1 and 2, 70 psi; curtain gas, 30 psi; temperature, 350°C; spray voltage, -4,500 V; declustering potential, -80 V; collision energy, -7 V; Qjet RF amplitude, 190 V. When the MRM^HR^ mode was used, the collision energy was set to -35 V.

### Nucleoside analysis by LC–MS/MS

For nucleoside analyses of in vitro methylated tRNAs, the Trm14 reaction was performed in 50 µL of buffer B containing 35 µM tRNA and 3 µM *T. kodakarensis* or *M. jannaschii* Trm14. The tRNA was purified by phenol extraction and precipitated by isopropanol. Recovered tRNA was dialyzed against water on a nitrocellulose membrane (0.025-μm VSWP, MF-Millipore, Merck) for 30 min. For nucleoside analyses of native tRNAs, isolated tRNAs were dialyzed against water on a nitrocellulose membrane. Forty picomoles of tRNA were digested by the following 3-step reactions at 37 °C for 1 h in each step: (I) 0.03 units of nuclease P1 (Fujifilm Wako Pure Chemical) in 10 mM NH_4_OAc (pH 5.3), (II) 0.04 units of phosphodiesterase I (Worthington Biochemical) in 50 mM ammonium bicarbonate (AMBIC) and (III) 0.03 units of alkaline phosphatase (*E. coli* C75, Nippon Gene) in 50 mM AMBIC. The samples were analyzed on an LC– MS system (UltiMate 3000 HPLC system and Q Exactive Orbitrap MS (Thermo Fisher Scientific)). Nucleosides were separated on a Hypersil GOLD aQ C18 LC column (150×2.1 mm, 1.9 µm, Thermo Fisher Scientific) with a guard cartridge column. The solvent containing 5 mM NH_4_OAc (pH 5.3) and acetonitrile (ACN) with a multi-step gradient (1–11% ACN from 0 to 6 min, 11–24% ACN from 6 to 12 min, 24–90% ACN from 12 to 15 min, and 90% ACN from 15 to 25 min at a flow rate of 100 μL/min), was used. The column was then re-equilibrated at 1% ACN from 25 to 34 min at 300 µL/min and 1% ACN from 34 to 35 min at 100 µL/min. Positive ion scanning ranged from 105 to 700 *m/z*. Mass spec data were analyzed using the Xcalibur Qual Browser (Thermo Fisher Scientific).

### Kinetic studies of *T. kodakarensis* Trm14 for SAM and tRNA

Kinetics was performed at 75 °C for 5 min in 10 µL of buffer C containing 50 mM Tris-HCl (pH 7.6), 50 mM KCl, 5 mM MgCl_2_, 6 mM 2-mercaptoethanol, 100 µM SAM, 100 nM [methyl-^3^H]-SAM, 3 µM Trm14, and 0.01-20 µM of *T. kodakarensis* tRNA^Cys^ transcript or 0.1-100 µM of *T. kodakarensis* tRNA^Trp^ transcript. For kinetic measurement of SAM, the SAM concentration was varied from 0.125 to 300 µM, and the tRNA concentration was increased to 20 µM.

### Gel assay of *T. kodakarensis* Trm14

For PAGE analysis, the Trm14 reaction was performed in 5 µL of buffer D containing 50 mM Tris-HCl (pH 7.6), 50 mM KCl, 5 mM MgCl_2_, 6 mM 2-mercaptoethanol, 35 µM tRNA, 100 µM [Methyl-^14^C]-SAM and 3 µM *T. kodakarensis* Trm14 at 75 °C for 10 min. The reaction was stopped by adding 15 µL of loading buffer containing 7 M urea, 1x TBE, and 0.1% bromophenol blue. Two microliters of the mixture were separated by 10% denaturing PAGE (7 M urea, 1x TBE). The gel was stained with methylene blue, destained with water, and dried. The incorporation of ^14^C-methyl groups in tRNAs was visualized with a phosphor imager.

### Disruption of *trm14* gene

The gene disruption plasmid pUTR14, previously constructed for deletion of the *trm14* gene (35), was used to transform the *T. kodakarensis* KUW1 strain, which exhibits uracil auxotrophy. Cells were grown in MA-YT-S⁰ medium at 85 °C for 10 h, harvested, and resuspended in 200 µL of 0.8× MA medium. After incubation on ice for 30 min, 3 µg of plasmid DNA was gently mixed with the cells and kept on ice for an additional 1 h. The transformation mixture was transferred to uracil-free ASW-AA-S⁰ liquid medium and incubated at 85 °C for 48 h. Subsequently, 200 µL of the culture was transferred to fresh medium and further incubated under the same conditions to enrich transformants displaying uracil prototrophy. Diluted cultures were spread onto ASW-AA solid medium and incubated at 85 °C for 3 days. Single colonies were isolated and cultivated, and genomic DNA was extracted using the phenol–chloroform method. Gene disruption was confirmed by PCR and DNA sequencing of the recombination region.

### Western blotting analysis of *T. kodakarensis* KUW1 and *Δtrm14* strain

*T. kodakarensis* cells were resuspended in a buffer containing 100 mM Tris-HCl (pH 6.8), 200 mM dithiothreitol, 2.5% SDS, 0.2% bromophenol blue, and 20% glycerol, and homogenized with an ultrasonic disruptor. Samples were boiled and immediately subjected to electrophoresis on a 12.5% SDS–polyacrylamide gradient gel. Proteins were transferred onto an Amersham Protran 0.45 NC membrane (Cytiva). *T. kodakarensis* Trm14 was detected using mouse antiserum containing anti-*T. kodakarensis* Trm14 polyclonal antibody (KITAYAMA LABES Co., Ltd.). Horseradish peroxidase (HRP) -conjugated Affinipure Goat Anti-Mouse IgG (H+L) (Proteintech) was used as a secondary antibody, and the chemiluminescence derived from HRP was detected with LuminoGraph EMI (ATTO).

### Characterization of the KUW1 (WT) and *Δtrm14* strains

To compare the growth phenotype of the KUW1 (WT) and *Δtrm14* strains, cells were grown in MA-YT-Pyr liquid medium. Pre-cultures were prepared by growing the strains in MA-YT-S⁰ medium for 12 h until the cells reached the stationary phase. Equal volumes of the pre-cultures were inoculated into fresh MA-YT-Pyr medium and incubated at 85 °C, 93 °C, or 95 °C. Cell growth was monitored by measuring the optical density at 660 nm (OD_660_) using a spectrophotometer.

### Phylogenetic analysis of Trm14- and TrmN-like proteins

Trm14-like proteins were identified by a BLASTP homology search against the UniProt protein database using *T. kodakarensis* Trm14 as the query sequence, which initially retrieved 250 candidate proteins. To exclude distantly related proteins such as Trm11, only 69 sequences with BLAST E-values < 1 × 10^−30^ were retained for the subsequent analyses. Multiple sequence alignment was performed using MAFFT version 7 (https://mafft.cbrc.jp/alignment/server/) with default parameters. A phylogenetic tree was constructed from the resulting alignment using the neighbor-joining (NJ) method with 100 bootstrap replicates and visualized using Archaeopteryx.js. Nineteen TrmN- like proteins were identified using the same procedure with *T. thermophilus* HB27 TrmN (TTC1157) as the query sequence. To compare the evolutionary relationships between Trm14- and TrmN-like proteins, the two sequence datasets were merged and analyzed by the same procedures described above.

## Results

### *Thermococcus kodakarensis* Trm14 possesses a dual-site specificity for G6 and G67

To determine whether Trm14 possesses a dual-site specificity for G6 and G67, we first obtained recombinant *T. kodakarensis* Trm14 with high purity through three purification steps (Fig. 2A). We also prepared *T. kodakarensis* tRNA^Cys^ and tRNA^Trp^ transcripts, which contain G6 and G67, respectively (Fig. 2B). We next assayed Trm14 activity and measured incorporation of the ^3^H-methyl group into the transcripts. We found that methylation of tRNA^Cys^ was more efficient than that of tRNA^Trp^ (Fig. 2C). Nucleoside analyses detected m^2^G (m/z 298.12, z = 1) in both reaction products (Fig. 2D). In contrast, m^2^_2_G (m/z 312.13, z = 1) was not detectable, indicating that m^2^G is the primary product of the Trm14 reaction, and thus the second methylation does not occur under the conditions tested, as reported previously (12). To identify the methylation sites, the methylated transcripts were digested with RNase A and analyzed by LC– MS/MS (Fig. 2E and 2F). For G67 analysis, *T. kodakarensis* tRNA^Gln^ transcript was used instead of *T. kodakarensis* tRNA^Trp^ transcript because RNase A digestion of the tRNA^Trp^ transcript does not generate a unique fragment containing G67, whereas the tRNA^Gln^ transcript yields the unique fragment 5′-GGGGCp-3′. For *T. kodakarensis* tRNA^Cys^ transcript, the fragment 5′-GGGAUp-3′ (m/z 843.109, z = –2) was detected in the absence of Trm14 (negative control) (Fig. 2E). In contrast, in the presence of Trm14, the relative abundance of this fragment decreased and a new fragment corresponding to 5′-GGGAUp-3′ + one methyl group (m/z 850.109, z = −2) appeared (Fig. 2E). MS/MS fragmentation patterns indicated that G6 in the tRNA^Cys^ transcript is methylated by Trm14 (Fig. 2F). Similarly, in the tRNA^Gln^ transcript, the fragment 5′-GGGGCp-3′ (m/z 850.613, z = –2) was detected in the absence of Trm14. In contrast, in the presence of Trm14, an RNA fragment 5′-GGGGCp-3′ + one methyl group (m/z 857.618, z = –2) appeared (Fig. 2G). MS/MS analyses demonstrated that G67 in the tRNA^Gln^ transcript was methylated (Fig. 2H). Kinetic analyses revealed differences in the apparent Km and Vmax values for the two tRNA substrates (Fig. 2I and 2J). *T. kodakarensis* Trm14 showed higher affinity for tRNA^Trp^ transcript with apparent Km value of 0.23 µM, which is approximately 6.5-fold higher affinity than that for tRNA^Cys^ transcript. In contrast, the Vmax value for the tRNA^Trp^ transcript was approximately 13-fold smaller than that for the tRNA^Cys^ transcript. The apparent Km for SAM was comparable among these two tRNA transcripts (Fig. 2I and 2J). These results show that *T. kodakarensis* Trm14 binds tRNA^Trp^ transcript with higher affinity but shows a lower turnover rate. This suggests that the sequence and/or structure of tRNA influence substrate recognition and reaction rates of Trm14. Overall, these biochemical data suggest that recombinant *T. kodakarensis* Trm14 catalyzes m^2^G formation at both positions 6 and 67 in tRNAs in vitro.

**Figure 2.**
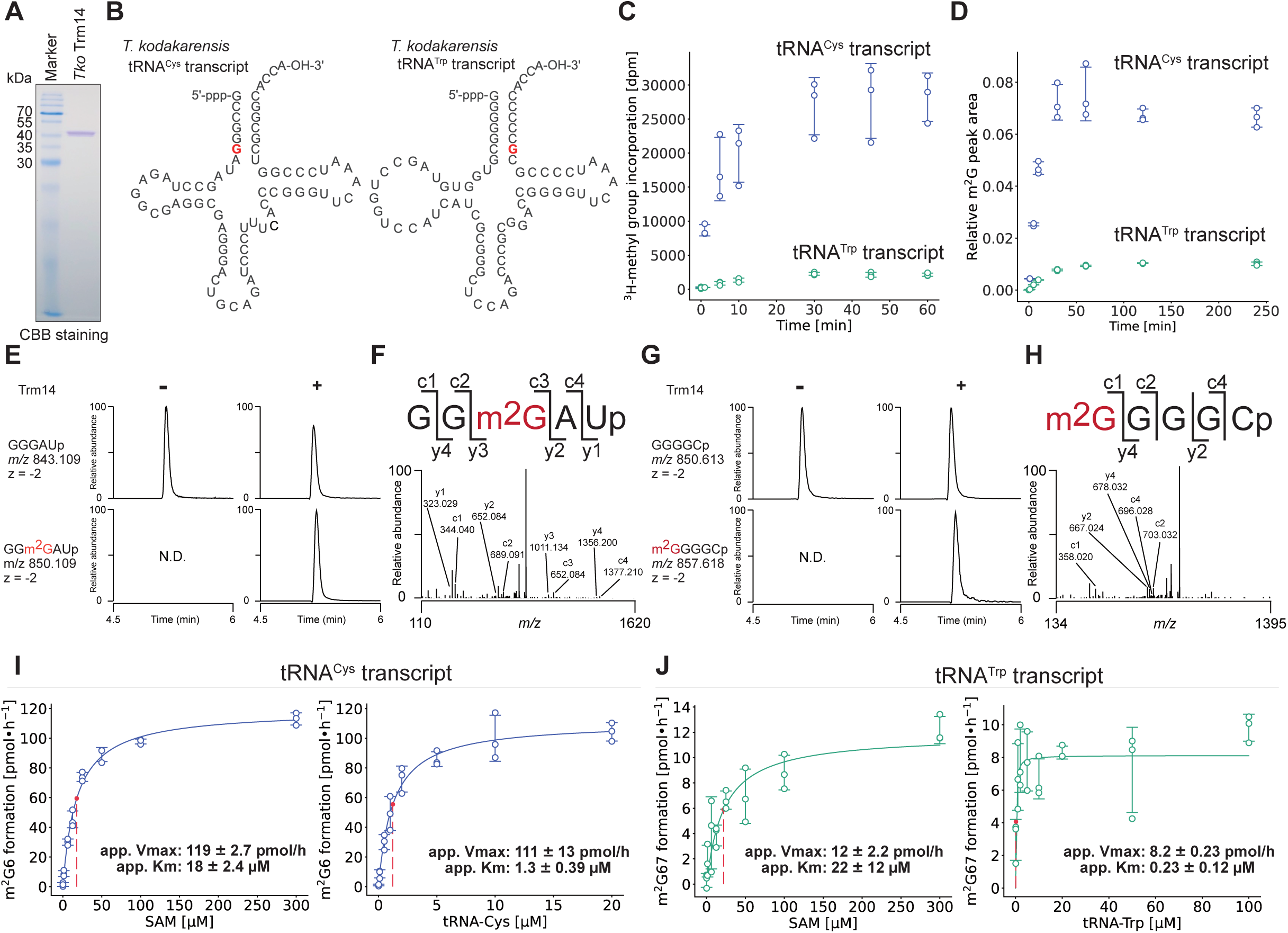
Recombinant *T. kodakarensis* Trm14 catalyzes m^2^G formation at positions 6 and 67 in tRNA. (A) Purified recombinant *T. kodakarensis* Trm14 (2 μg) was analyzed by 15% SDS- PAGE and visualized by Coomassie Brilliant Blue (CBB) staining. (B) Cloverleaf secondary structures of *T. kodakarensis* tRNA^Cys^ transcript (left) as well as tRNA^Trp^ transcript (right) are depicted. G6 and G67 are highlighted in red. (C) Methyltransferase activity of recombinant *T. kodakarensis* Trm14 was measured at 75 °C using tRNA^Cys^ (blue) and tRNA^Trp^ (green) transcripts in the presence of ^3^H-labeled SAM. Experiments were performed independently three times (n = 3). Error bars indicate standard deviations. (D) The m^2^G levels in methylated tRNA transcripts were quantified by nucleoside analyses. Data from tRNA^Cys^ and tRNA^Trp^ transcripts are indicated by blue and green circles, respectively. The relative abundance of m^2^G was normalized to that of cytidine. (E–H) Methylation sites in (E, F) *T. kodakarensis* tRNA^Cys^ and (G, H) tRNA^Gln-CTG^ transcripts were determined by LC–MS/MS. Extracted ion chromatograms corresponding to RNase A-digested RNA fragments containing (E) m^2^G6 or (G) m^2^G67 are shown. CID spectra of the (F) m^2^G6-containing and (H) m^2^G67-containing fragments of tRNA^Cys^ and tRNA^Gln^ transcripts, respectively. N.D. stands for Not Detected. (I and J) Kinetic analyses of recombinant *T. kodakarensis* Trm14 using (I) tRNA^Cys^ and (J) tRNA^Trp^ transcripts. Apparent kinetic parameters for SAM (left panels) and tRNA substrates (right panels) were determined. Experiments were performed independently three times (n = 3). Error bars indicate standard deviations. Data were fitted to the Michaelis-Menten equation.

We next constructed a *trm14* gene disruptant of *T. kodakarensis* by homologous recombination (Supplementary Fig. 1A). Disruption of the *trm14* gene was confirmed by polymerase chain reaction (Supplementary Fig. 1B and 1C) and western blotting analysis using an anti-*T. kodakarensis* Trm14 polyclonal antibody (Fig. 3A). Two bands were detected at approximately 40 kDa in the KUW1 (WT) strain, suggesting cross-reactivity of the polyclonal antibody. The upper band disappeared in the Δ*trm14* strain, and recombinant Trm14 migrated at the same position, indicating that the upper band corresponds to Trm14. We hypothesized that the lower band originated from another THUMP-domain-containing protein because this RNA-binding domain is conserved among multiple tRNA modification enzymes. One candidate was TrmTS, which also contains a THUMP domain and catalyzes 2’-*O*-methylation of cytidine at position 6 in *T. kodakarensis* tRNA^Trp^ (36). To test this, western blotting analyses were performed using a cell extract from a Δ*trmTS* strain (36) and purified recombinant TrmTS protein. As expected, the lower band disappeared in the Δ*trmTS* strain, and recombinant TrmTS migrated at the same position as the lower band detected in the WT strain. Together, these results confirmed successful disruption of the *trm14* gene.

**Figure 3.**
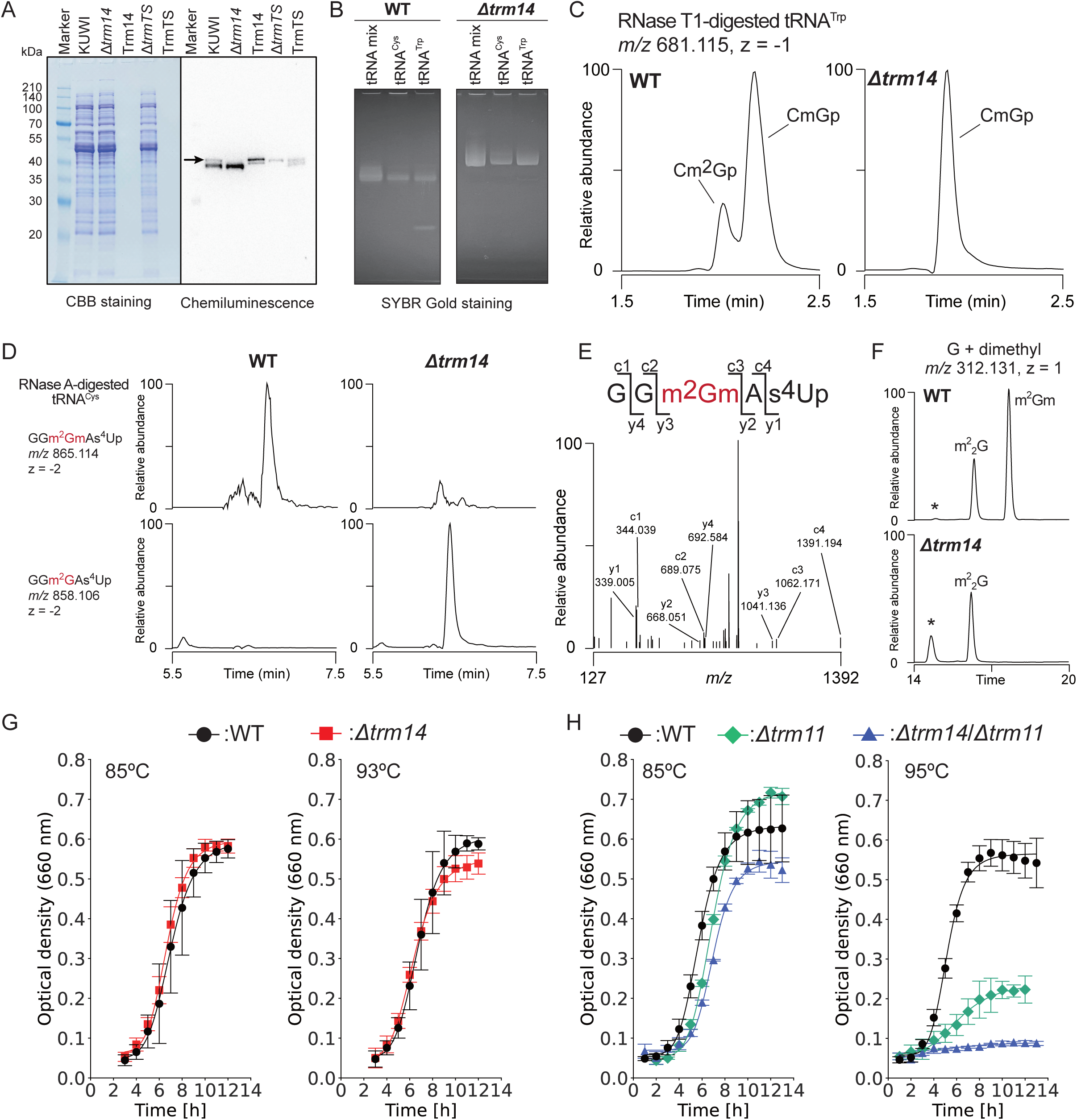
*Thermococcus kodakarensis* Trm14 catalyzes m^2^G formation at both G6 and G67 in vivo. (A) Western blotting analyses of Trm14 were performed using cell extracts from the WT (KUW1), *Δtrm14*, and *ΔtrmTS* strains. Purified recombinant Trm14 and TrmTS proteins were included as migration controls. Because only 20 ng of recombinant Trm14 and 10 ng of recombinant TrmTS were loaded, these proteins were not detectable by CBB staining. The Trm14 band is indicated by a black arrow. (B) Native tRNA^Cys^ and tRNA^Trp^ isolated from WT and *Δtrm14* strains were analyzed by 10% denaturing PAGE (7 M urea). (C) Extracted ion chromatograms of the indicated negative ions from RNase T1-digested tRNA^Trp^ from WT and Δ*trm14* strains are shown. CID spectra of the observed peaks are provided in Supplementary Fig. 2. (D) Extracted ion chromatograms of the indicated negative ions from RNase A-digested tRNA^Cys^ from WT and Δ*trm14* strains are shown. (E) CID spectrum of the m^2^Gm-containing fragment of tRNA^Cys^. (F) Extracted ion chromatograms of the indicated positive ions corresponding to guanosine, mono-methylated guanosine, and di-methylated guanosine are shown. The asterisk indicates an unidentified peak. (G) Growth curves of the WT (black) and Δ*trm14* (red) strains cultured at 85 °C and 93 °C. Data are presented as the mean of independent biological replicates (n = 9 and n = 11 for WT and Δ*trm14* strain, respectively). Error bars indicate standard deviations. (H) Growth curves of the WT (black), Δ*trm11*, and Δ*trm11*-Δ*trm14* strains cultured at 85 °C and 95 °C. Data are presented as an average of independent biological replicates (n =6 or n = 3). Error bars indicate standard deviations.

Native *T. kodakarensis* tRNA^Cys^ and tRNA^Trp^ were purified from the WT and Δ*trm14* strains using a solid-phase DNA probe method (41,42) (Fig. 3B). The purified tRNAs were digested with RNase T1 or RNase A and analyzed by LC–MS/MS (Fig. 3C-3F). For the native tRNA^Trp^, RNase T1 digestion generated two methylated fragments, 5′-Cm^2^Gp-3′ and 5′-CmGp-3′, in the WT strain (Fig. 3C and Supplementary Fig. 2A), consistent with our previous study (35). Notably, the first peak was absent in the tRNA^Trp^ purified from the Δ*trm14* strain (Fig. 3C and Supplementary Fig. 2B), demonstrating that Trm14 catalyzes m^2^G67 formation in vivo. For the native tRNA^Cys^, intact tRNA was first analyzed by LC–MS. The observed molecular masses were 24,726.86 Da for the WT tRNA^Cys^ and 24,712.90 Da for the Δ*trm14* tRNA^Cys^, corresponding to a mass difference of 14 Da, equivalent to a single methyl group (Supplementary Fig. 3). RNase A digestion of WT tRNA^Cys^ generated a fragment corresponding to 5′-GGGAU-3′ + two methyl groups and a sulfur modification (m/z 865.114, z = –2) (Fig. 3D). In contrast, the observed m/z for the corresponding fragment from the Δ*trm14* strain shifted to 858.106 (z = –2) (Fig. 3D), indicating the loss of one methyl group. MS/MS analysis of this fragment derived from the WT tRNA^Cys^ revealed that the two methyl groups were installed on G6 (Fig. 3E). We also assigned the sulfur modification detected in this fragment as s^4^U8 (Fig. 3E). In addition, nucleoside analyses were performed to determine which guanosine modification was affected by disruption of the *trm14* gene (Fig. 3F and Supplementary Fig. 4A-4I). A peak corresponding to m^2^Gm was detected in WT tRNA^Cys^, whereas the Δ*trm14* tRNA^Cys^ showed complete loss of the m^2^Gm peak (Fig. 3F). Each peak observed in the extracted ion chromatograms of G (m/z 284.010, z = 1), mono-methylated G (m/z 298.115, z = 1), and di-methylated G (m/z 312.131, z = 1) was further analyzed by MS/MS (Supplementary Fig. 4A-4I). Comparison of MS/MS fragmentation patterns for mono- methylated G revealed that the relative abundance of Gm (m/z 152.057, a product ion corresponding to a guanine base, z = 1) remarkably increased in the Δ*trm14* strain (Supplementary Fig. 4C and 4G), consistent with the loss of m^2^Gm in the Δ*trm14* strain (Fig. 3F). An unidentified peak with m/z 312.131 (z = 1), corresponding to di- methylated G, also increased in the Δ*trm14* strain. Overall, these analyses indicate that Trm14 catalyzes m^2^G formation in the fragment 5′-GGm^2^GmAs^4^Up-3′. Together, the intact tRNA, RNA fragment, and nucleoside MS analyses demonstrate that Trm14 catalyzes m^2^G formation at both positions 6 and 67 in tRNAs in vivo.

**Figure 4.**
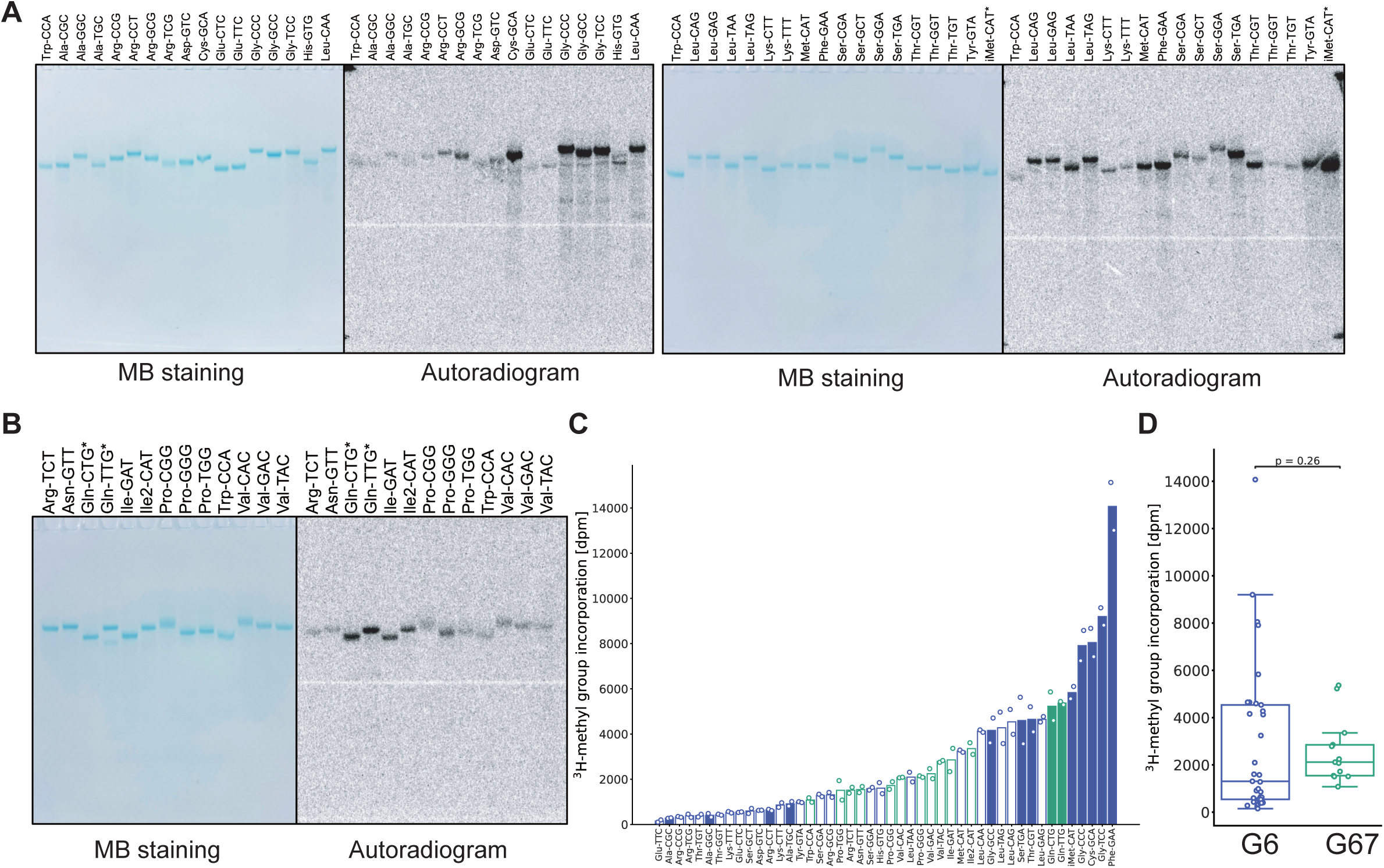
*T. kodakarensis* Trm14 methylated all *T. kodakarensis* tRNA transcripts in vitro. (A and B) *T. kodakarensis* tRNA transcripts containing (A) G6 or (B) G67 were reacted with recombinant *T. kodakarensis* Trm14 in the presence of ^14^C-labeled SAM. Methylated tRNAs were analyzed by 10% denaturing PAGE (7 M urea), and incorporation of ^14^C-methyl group in tRNAs was visualized by autoradiography. *T. kodakarensis* tRNA^Trp^ transcript was included as a control on each gel. Asterisks show tRNAs starting from non-G. These tRNAs were prepared as a precursor tRNA transcript (i.e. a leader sequence-containing tRNA transcript), and then the 5’-leader sequence was removed by RNase P. (C) Methyl-group incorporation into individual tRNA transcripts after a 10-min reaction at 75 °C. Each tRNA transcript was analyzed in two independent experiments (n = 2). Transcripts containing G6 and G67 are indicated by blue and green bars, respectively. Filled bars denote tRNAs containing an A-U or G-U base pair within the acceptor stem. (D) Summary of methyl-group incorporation into G6-containing and G67-containing tRNA transcripts. Statistical significance was assessed using the Mann– Whitney U test.

To analyze physiological roles of these modified nucleosides, we next examined the growth phenotype of the Δ*trm14* strain at 85 °C and 93 °C. We found that its growth rate was comparable to that of the WT strain (Fig. 3G). We also analyzed a Δ*trm14/*Δ*trm11* double disruptant strain constructed in our previous study (35). In this strain, both m^2^G67 and m^2^_2_G10 are absent from *T. kodakarensis* tRNA^Trp^ (35). As reported previously, the Δ*trm11* strain showed growth retardation at high temperatures (Fig. 3H). The Δ*trm14*/Δ*trm11* strain exhibited more severe growth retardation than the Δ*trm11* strain at high temperatures (Fig. 3H). Thus, these results indicate that m^2^G6/m^2^G67 function cooperatively with m^2^G10/m^2^_2_G10 to support cellular fitness at high temperatures.

Overall, these analyses revealed the loss of m^2^G6 in tRNA^Cys^ and m^2^G67 in tRNA^Trp^ in the Δ*trm14* strain, and therefore Trm14 is responsible for m^2^G formation at both positions in vivo and in vitro. In addition, the results from in vitro experiments with the purified Trm14 demonstrated that neither other subunit proteins nor guide RNAs are necessary for the Trm14 catalysis, and thus Trm14 is a stand-alone enzyme.

### *Thermococcus kodakarensis* Trm14 methylates all tRNA species

The biochemical and genetic analyses described above demonstrated that Trm14 catalyzes m^2^G formation at both G6 and G67. According to the genomic tRNA database (43), 46 distinct tRNA genes are encoded in the *T. kodakarensis* genome. We noticed that all tRNAs contain either G6 or G67 in their sequences. Based on the dual-site specificity of Trm14, we hypothesized that all *T. kodakarensis* tRNAs could be substrate tRNAs for this enzyme. To test this, we prepared all 46 tRNA transcripts by in vitro transcription, performed Trm14 reactions at 75 °C for 10 min, and measured ^14^C-methyl-group incorporation into each transcript (Fig. 4). Gel-based assays demonstrated that all tested tRNA transcripts are methylated by Trm14, although the band intensities varied among tRNA species (Fig. 4A and 4B). To quantify these differences more accurately, we performed a conventional filter assay and measured ^3^H-methyl group incorporation into all transcripts (Fig. 4C and 4D). Among G6-containing tRNAs, tRNA^Phe-GAA^ showed the highest methyl group incorporation activity, whereas tRNA^Glu- TTC^ and tRNA^Ala-CGC^ exhibited approximately 90 and 50-fold decreased activities, respectively. Similarly, among G67-containing tRNAs, tRNA^Gln-TTG^ was efficiently methylated, whereas tRNA^Trp-CCA^ showed approximately 5-fold decreased methyl group incorporation. Despite these tRNA-dependent differences, statistical analysis revealed no significant difference in methyl group incorporation between the G6- and G67-tRNA species (Fig. 4D). These results suggest that Trm14 broadly recognizes all tRNA species in *T. kodakarensis* and that methylation efficiency is influenced more by individual tRNA sequence than by the target position itself.

We next sought to determine what factors govern the methylation efficiency of individual tRNAs. Based on the genomic tRNA sequences of *T. kodakarensis*, we noticed that highly methylated tRNAs frequently contain an A-U or G-U pair adjacent to the target site in the acceptor stem (Fig. 4C, filled bars). Among *T. kodakarensis* tRNAs, four distinct A-U/G-U positions (1–72, 3–70, 5–68, and 7–66) are found within the acceptor stem. Interestingly, tRNAs containing A3-U70 or G3-U70, including tRNA^Ala-CGC^, tRNA^Ala-GGC^, tRNA^Ala-TGC^, and tRNA^Arg-CCT^, were methylated by Trm14 with relatively low methylation levels despite the presence of an A-U base pair (Fig. 4C, filled bars). In contrast, the remaining A-U/G-U pairs are located either adjacent to the target site or at the end of the acceptor stem and were generally methylated with higher methylation levels (Fig. 4C, filled bars). Thus, these observations suggest that the position of the A-U/G-U base pair, rather than its mere presence, might influence Trm14 activity. Because base-pair stability is determined by both hydrogen-bonding and stacking interactions with a neighboring base pair, we checked the thermodynamic stability of representative nearest-neighbor base pairs. The ΔG°_37_ values of 5′- r(CG/GC)-3′, 5′-r(AC/UG)-3′ (internal stack), 5′-r(CA/GU)-3′ (helix end), and 5′- r(CG/GU)-3′ (helix end) are −2.33, −2.25, −1.63, and −0.92 kcal/mol, respectively (44–46). Interestingly, the methyl group incorporation levels tended to decrease with increasing thermodynamic stability. Because the A-U/G-U base pairs at positions 3–70 are 3 nt away from the target sites, they are not expected to affect the thermodynamic stability of the target-site, according to the nearest-neighbor model (44). Therefore, we hypothesized that local thermodynamic stability adjacent to the target site might influence Trm14 activity. To test this, we prepared two groups of mutant tRNA transcripts: one with weakened acceptor stems and the other with strengthened acceptor stems. First, we prepared four tRNA^Leu^ mutant transcripts in which the G-C base pair at positions 1–72, 3–70, 5–68, or 7–66 was individually replaced with an A-U or G-U base pair (Fig. 5A). Trm14 assays showed increased methyl group incorporation into the tRNA^Leu^ (G1A, C72U), (G5U, C68A), (G7A, C66U), and (C66U) mutants, whereas no statistically significant enhancement was observed for the tRNA^Leu^ (G3A, C70U) mutant (Fig. 5B). We next prepared three mutant transcripts, tRNA^Cys^ (A7G, U66C), tRNA^Phe^ (U66C), and tRNA^Gln^ (A1G, U72C), in which the acceptor stem was strengthened (Fig. 5C–E). In the case of tRNA^Cys^ (A7G, U66C) and tRNA^Phe^ U66C mutant transcripts, strengthening the acceptor stem significantly reduced methyl group incorporation by Trm14 (Fig. 5F and 5G). In contrast, while the average value of relative methyl group incorporation into the tRNA^Gln^ (A1G, U72C) mutant transcript decreased as compared with that into the tRNA^Gln^ transcript, the difference was not statistically significant (Fig. 5H). It should be noted that all assays were performed at 75°C. Together, these results indicate that the local stability of the acceptor stem surrounding the target site is a major determinant of Trm14 activity.

**Figure 5.**
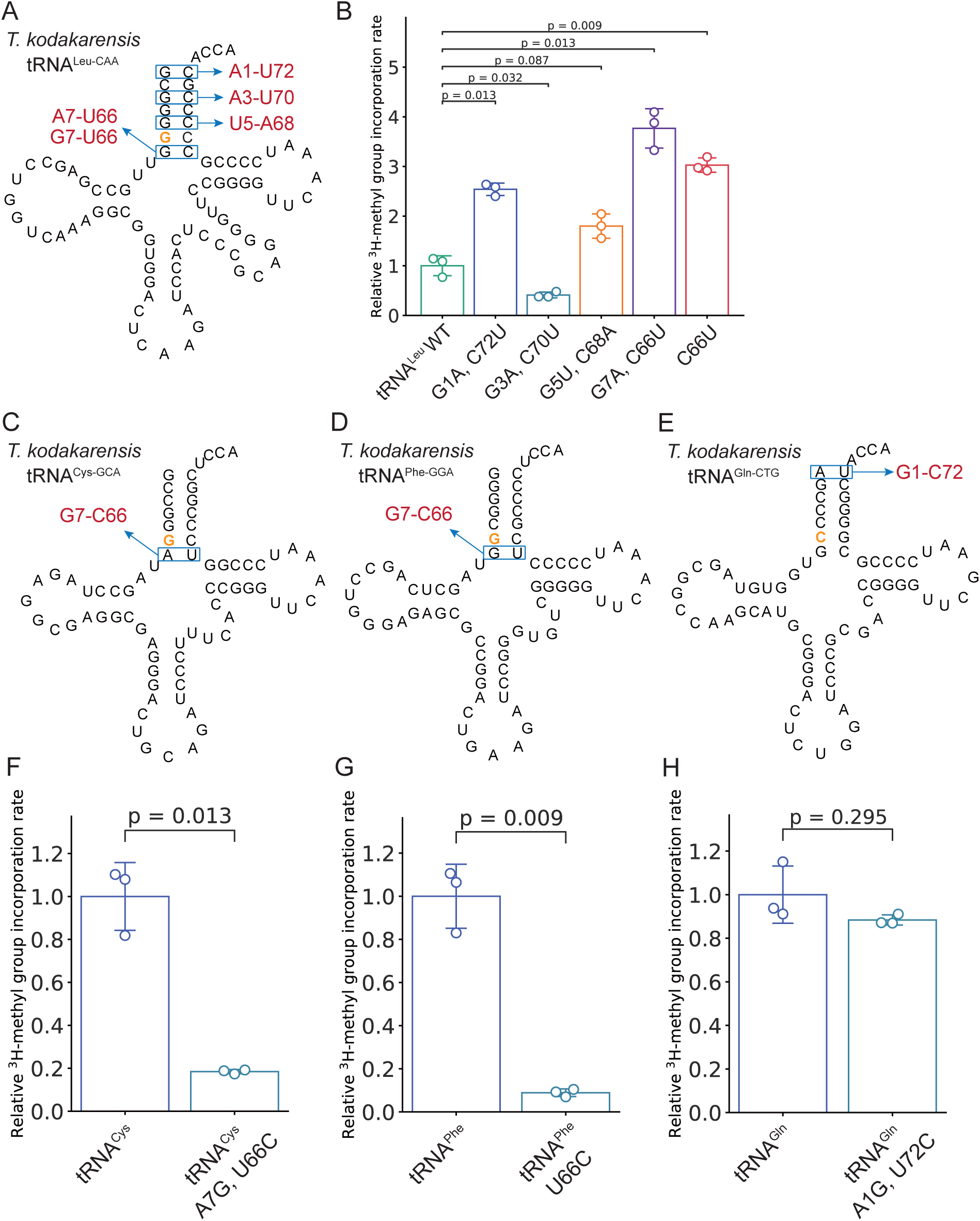
The local stability of the acceptor stem surrounding the target site regulates methylation by *T. kodakarensis* Trm14. (A) Mutant tRNA^Leu-CAA^ transcripts in which a G-C base pair in the acceptor stem was replaced with an A-U or G-U base pair, are depicted. (B) ^3^H-methyl group incorporation into WT and mutant tRNA^Leu^ transcripts after a 10-min reaction at 75 °C was quantified. Experiments were performed independently three times (n = 3). Statistical significance was assessed using Student’s t-test. Error bars indicate standard deviations. (C-E) An A- U or G-U base pair in the acceptor stem in (C) tRNA^Cys-GCA^, (D) tRNA^Phe-GGA^, and (E) tRNA^Gln-CTG^ was substituted with a G-C pair. (F-H) ^3^H-methyl group incorporation into WT and mutant (F) tRNA^Cys-GCA^, (G) tRNA^Phe-GGA^, and (H) tRNA^Gln-CTG^ transcripts after a 10 min reaction at 75 °C was quantified. Experiments were performed independently three times (n = 3). Statistical significance was assessed using Student’s t-test. Error bars indicate standard deviations.

### Disruption of the G6-C67/C6-G67 base pair enhances methylation by *T. kodakarensis* Trm14

Based on the biochemical analyses, we hypothesized that the base-pairing status of G6- C67 and C6-G67 could influence methylation efficiency by Trm14. To gain further mechanistic insight into substrate recognition and catalysis by archaeal Trm14, we prepared mutant tRNA^Cys^ (C67A) and tRNA^Trp^ (C6A) transcripts in which base pairs at the target sites were disrupted and examined their methyl group incorporation activity using *T. kodakarensis* Trm14 (Fig. 6A-6D). Interestingly, methyl group incorporation into both mutant transcripts was significantly increased relative to the corresponding WT transcripts (Fig. 6B). At the 10-min time point, methyl group incorporation into the tRNA^Cys^ C67A and tRNA^Trp^ C6A mutant transcripts was approximately 2-fold and 5- fold higher, respectively, than that observed for the corresponding WT transcripts (Fig. 6C and 6D). To examine the reaction products, we performed nucleoside analyses of the methylated transcripts. In both mutant tRNAs, only m^2^G was detected, whereas m^2^_2_G remained not detectable (Fig. 6E). Together, these results indicate that disruption of the base pairs at the target sites enhances methylation by Trm14 but does not promote m^2^_2_G formation at least under the tested conditions. Thus, disruption of base pairing accelerates the Trm14 reaction while maintaining its strict specificity for mono- methylation. In addition, Trm14 does not require a base pair at the target site for methylation.

**Figure 6.**
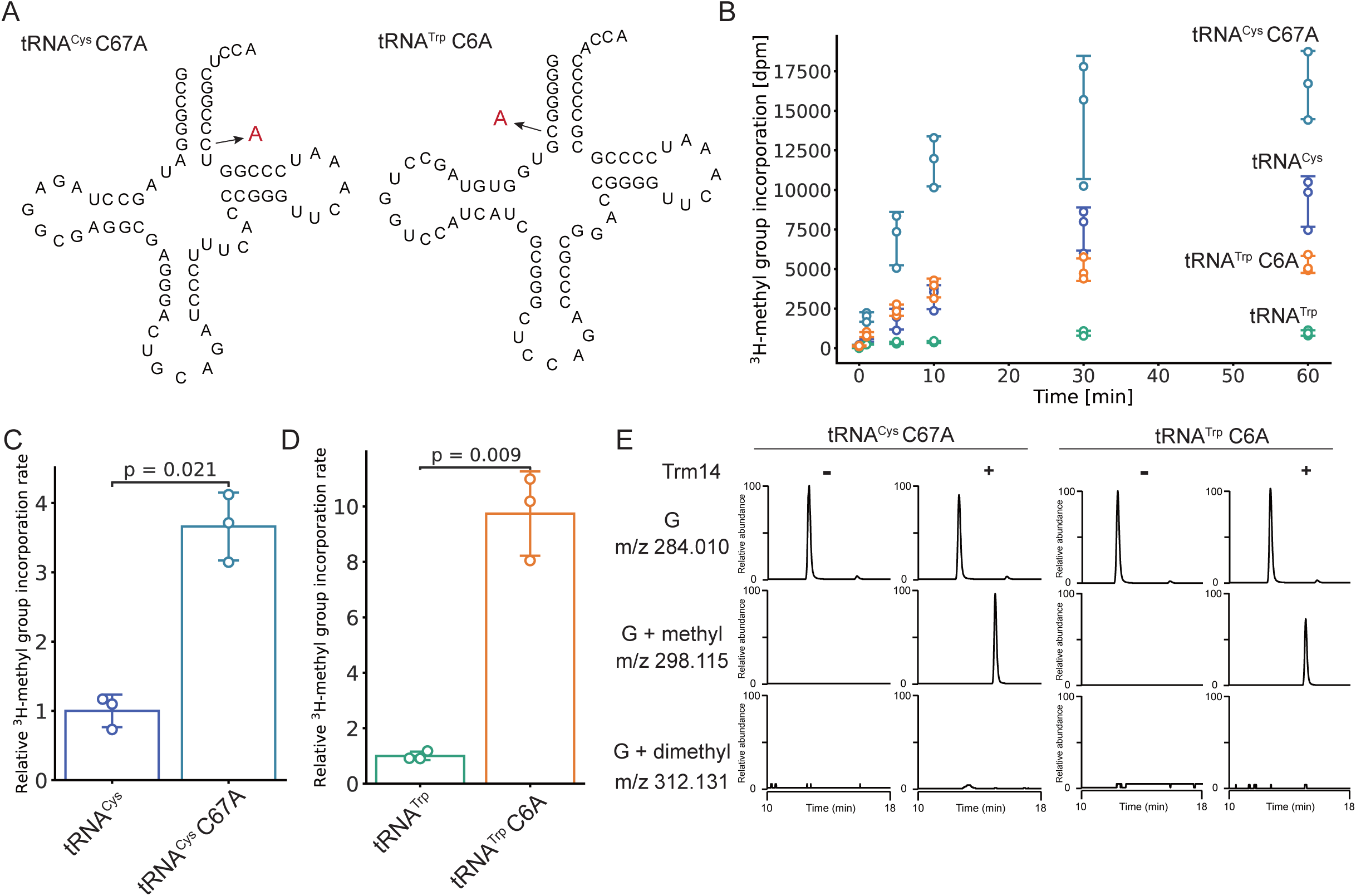
Disruption of the target-site base pair enhances methylation by *T. kodakarensis* Trm14. (A) Cloverleaf structures of mutant tRNA^Cys^ and tRNA^Trp^ transcripts are shown. In the mutant transcripts, C67 of tRNA^Cys^ and C6 of tRNA^Trp^ were individually substituted with A to disrupt the target-site base pair. (B) Time-course analysis of ^3^H-methyl-group incorporation into tRNA^Cys^, tRNA^Trp^, tRNA^Cys^ C67A, and tRNA^Trp^ C6A transcripts by recombinant *T. kodakarensis* Trm14 was performed at 75 °C. (C and D) ^3^H-methyl-group incorporation into (C) tRNA^Cys^ and tRNA^Cys^ C67A transcripts and (D) tRNA^Trp^ and tRNA^Trp^ C6A transcripts after a 10-min reaction at 75 °C was quantified. Experiments were performed independently three times (n = 3). Statistical significance was assessed using Student’s t-test. Error bars indicate standard deviations. (E) Nucleoside analyses of methylated tRNA^Cys^ C67A and tRNA^Trp^ C6A transcripts by LC–MS/MS. Extracted ion chromatograms (XICs) corresponding to G, mono-methylated G (m^2^G), and di-methylated G (m^2^_2_G) are shown.

Dual-site specificity is conserved in *M. jannaschii* Trm14.

Trm14 was originally identified as a tRNA m^2^G6 methyltransferase in *M. jannaschii* (12). In the earlier study, *M. jannaschii* tRNA^Cys^ (G6), tRNA^Pro^ (U6 for negative control), and tRNA^Asp^ (G6) transcripts were tested for *M. jannaschii* Trm14 activity, whereas no G67-containing tRNA substrates were examined (12). We therefore asked whether the dual-site specificity observed for *T. kodakarensis* Trm14 is conserved in other archaeal Trm14 enzymes. To test this, we prepared recombinant *M. jannaschii* Trm14 (Fig. 7A) and performed in vitro methylation assays using *M. jannaschii* tRNA^Cys^ (G6) and tRNA^Ile2^ (G67) transcripts. *M. jannaschii* Trm14 efficiently methylated both substrates (Fig. 7B). We also tested *T. kodakarensis* tRNA^Cys^ (G6) and tRNA^Trp^ (G67) transcripts as substrates for *M. jannaschii* Trm14. Interestingly, *M. jannaschii* Trm14 showed substrate preferences similar to those of *T. kodakarensis* Trm14, showing higher methylation activity toward tRNA^Cys^ transcript than toward tRNA^Trp^ transcript (Fig. 7B). To identify the methylation site in *M. jannaschii* tRNA^Ile2^, the methylated transcript was digested with RNase A and analyzed by LC–MS/MS (Fig. 7C and 7D). An RNA fragment corresponding to 5′-GGGCp-3′ + one methyl group (m/z 685.095, z = –2) was detected after Trm14 reaction (Fig. 7C). Furthermore, MS/MS analysis demonstrated that the methyl group was installed on G67 in tRNA^Ile2^ (Fig. 7D). Similarly, in the case of *M. jannaschii* tRNA^Cys^ transcript, an RNA fragment corresponding to 5′-GGGUp-3′ + one methyl group (m/z 571.738, z = –3) was detected after Trm14 reaction (Supplementary Fig. 5A). Also, MS/MS analysis demonstrated that the methyl group was installed on G6 in tRNA^Cys^ (Supplementary Fig. 5B), as reported previously (12). Together, we concluded that *M. jannaschii* Trm14 also catalyzes methylation at both G6 and G67. These results indicate that dual-site specificity is conserved at least in these two different archaeal Trm14 enzymes.

**Figure 7.**
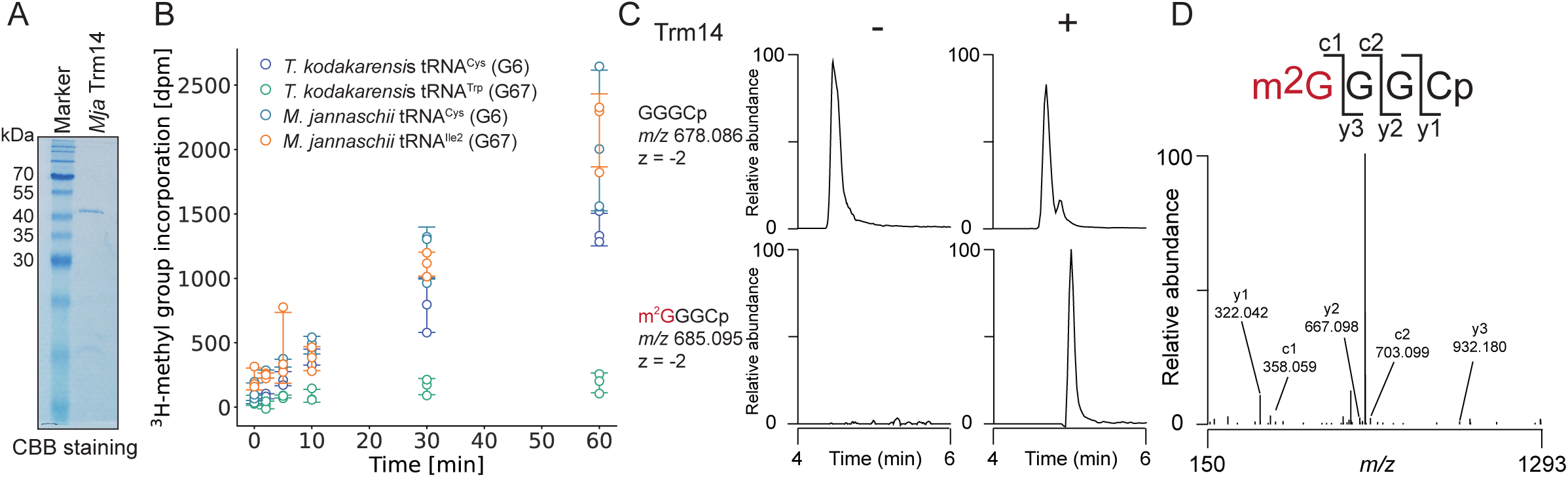
Recombinant *M. jannaschii* Trm14 catalyzes methylation at both G6 and G67. (A) Purified recombinant *M. jannaschii* Trm14 (2 μg) was analyzed by 15% SDS-PAGE and visualized by CBB staining. (B) Methyl group transfer activity of recombinant *M. jannaschii* Trm14 toward *T. kodakarensis* tRNA^Cys^ and tRNA^Trp^ transcripts, and *M. jannaschii* tRNA^Cys^ and tRNA^Ile2^ transcripts. ^3^H-methyl group incorporation was monitored over time. Experiments were performed independently three times (n = 3). Error bars indicate standard deviations. (C) Extracted ion chromatograms of the indicated negative ions from RNase A-digested *M. jannaschii* tRNA^Ile2^ transcripts with or without recombinant *M. jannaschii* Trm14 are shown. (D) CID spectrum of the m^2^G- containing fragment of tRNA^Ile2^.

### Dual-site specificity is not conserved in TrmN, a bacterial ortholog of Trm14

TrmN is the bacterial ortholog of Trm14 and catalyzes methylation of guanosine at position 6 in tRNAs from *T. thermophilus* (15). Trm14 and TrmN share a similar domain architecture, and their overall structures are highly similar (17). To determine whether TrmN, like *T. kodakarensis* Trm14, can methylate both G6 and G67 in tRNAs, we prepared recombinant *T. thermophilus* TrmN (Fig. 8A) and examined its activity for several tRNA transcripts. In addition to tRNA^Phe^, which is currently the only reported *T. thermophilus* tRNA containing m^2^G6 (6), we prepared tRNA^Tyr^ (G6), tRNA^Thr^ (U6), tRNA^Asp^ (G67), and tRNA^Glu^ (G67) transcripts (Fig. 8B). Consistent with our previous study, recombinant TrmN methylated the tRNA^Phe^ transcript. Furthermore, TrmN also methylated tRNA^Tyr^ (G6) but showed no detectable activity for tRNA^Thr^ (U6), tRNA^Asp^ (G67), or tRNA^Glu^ (G67). These results demonstrate that *T. thermophilus* TrmN specifically recognizes G6-containing tRNAs and lacks the dual-site specificity observed for *T. kodakarensis* Trm14. Thus, despite their overall structural similarity, TrmN and Trm14 appear to employ distinct mechanisms for substrate recognition.

**Figure 8.**
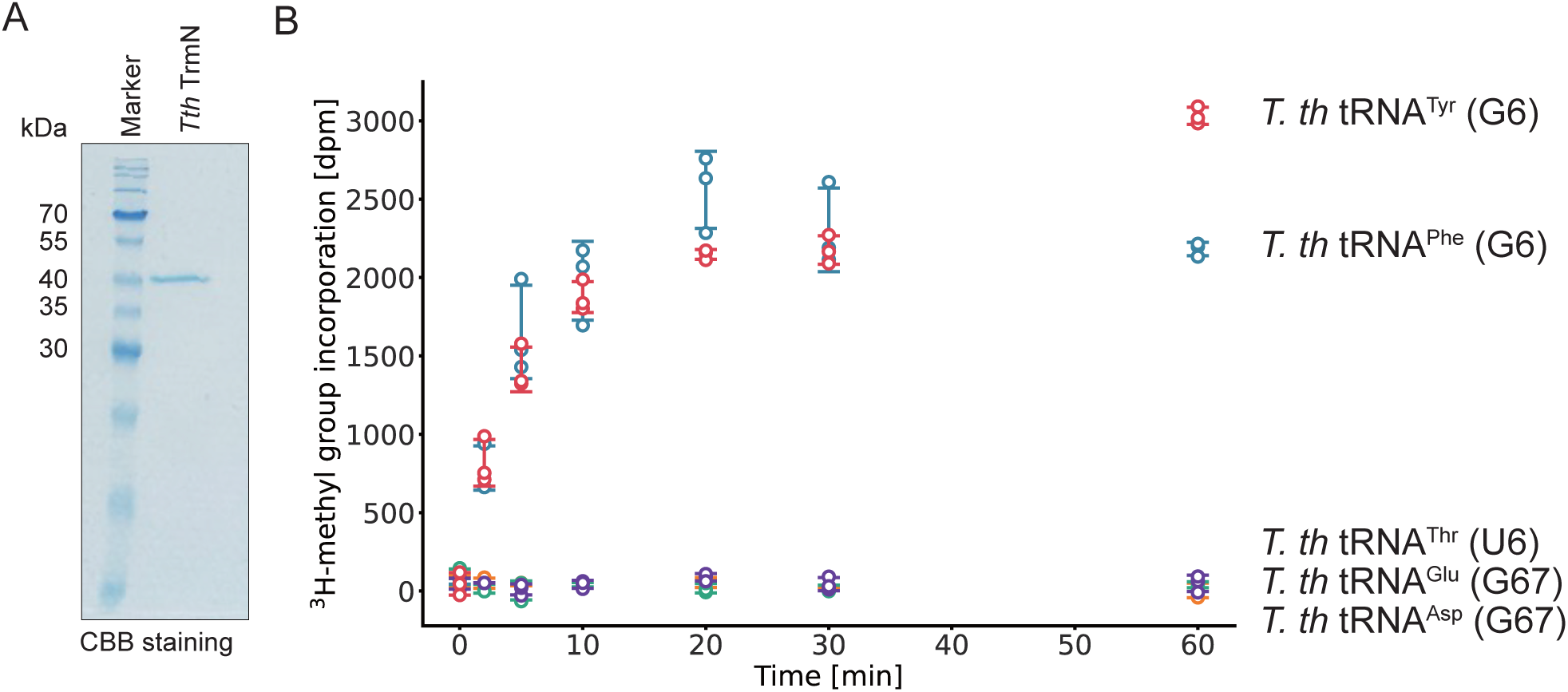
*T. thermophilus* TrmN specifically methylates G6-containing tRNAs. (A) Purified recombinant *T. thermophilus* TrmN (2 μg) was analyzed by 15% SDS- PAGE and visualized by CBB staining. (B) Methyl-group transfer activity of recombinant *T. thermophilus* TrmN toward *T. thermophilus* tRNA^Tyr^ (G6), tRNA^Phe^ (G6), tRNA^Thr^ (U6), tRNA^Glu^ (G67), and tRNA^Asp^ (G67) transcripts. ^3^H-methyl-group incorporation was monitored over time. Experiments were performed independently three times (n = 3). Error bars indicate standard deviations.

## Discussion

Although numerous modified nucleosides have been identified in tRNAs, the enzymes responsible for many of these modifications remain unidentified or have been proposed but not experimentally validated (4). In our previous study, we determined the primary sequence of *T. kodakarensis* tRNA^Trp^ and identified 21 modified nucleosides and predicted the enzymes responsible for these modifications (35). However, only a limited number of these enzymes, including archaeal Trm11, ThiI, TYW1, and TrmTS, have been experimentally characterized in *T. kodakarensis* (35,36). To bridge this gap, we focused on archaeal Trm14 and comprehensively characterized its biochemical and genetic properties in the present study. We found that both *T. kodakarensis* Trm14 and *M. jannaschii* Trm14 exhibit dual-site specificity toward G6 and G67 in tRNA. Because only two Trm14 enzymes were experimentally characterized in the current study, we examined how broadly this property might be conserved among archaeal Trm14 proteins. Phylogenetic analysis of Trm14-like proteins showed that *T. kodakarensis* and *M. jannaschii* Trm14 belong to evolutionarily distinct branches within Euryarchaeota (Supplementary Fig. 6), suggesting that dual-site specificity is conserved at least among euryarchaeal Trm14 enzymes. A separate phylogenetic analysis including both Trm14- and TrmN-like proteins showed that archaeal Trm14 homologs form a clade distinct from the predominantly bacterial TrmN clade, although a small group of proteins was identified by both Trm14- and TrmN-based homology searches (Supplementary Fig. 7; highlighted in green). This overlap indicates that these proteins share sequence similarity with both enzyme groups despite their phylogenetic separation. These proteins may have a different enzymatic property from Trm14 or TrmN.

Most tRNA modifications outside the anticodon loop are individually dispensable for cellular growth, as disruption of a tRNA modification enzyme often shows no detectable phenotype. This apparent dispensability is generally attributed to functional redundancy among multiple modifications that collectively stabilize tRNA. For example, m^2^G10 and m^2^_2_G26 are closely located and stacked within the tertiary core of yeast tRNA^Phe^, where they cooperatively stabilize the tRNA structure (32). Although deletion of either the *trm11* or *trm1* gene alone causes little growth defect, simultaneous disruption of both genes results in severe growth retardation (47). Similarly, simultaneous loss of the tRNA m^7^G46 methyltransferase Trm8 and the tRNA m^5^C49 methyltransferase Trm4 activates rapid tRNA decay and causes a severe growth defect in yeast (48,49), whereas depletion of the corresponding human enzymes increases cellular sensitivity to 5-fluorouracil (50). Additional examples, including the combined loss of s^4^U8 and m^5^U54 or of ac^4^C12 and Um44, likewise demonstrate that multiple modifications cooperate to maintain tRNA stability and prevent tRNA degradation (51–53). Loss of these modifications can also impair aminoacylation efficiency and alter gene expression through defects in translation (54). Notably, many of these functionally redundant modifications are clustered within the elbow region of tRNA, where they stabilize the tertiary structure. Our genetic analyses revealed the physiological significance of Trm14 and m^2^G6/m^2^G67 in *T. kodakarensis*. The growth phenotype of the Δ*trm14* strain was comparable to that of the WT strain at high temperatures. In contrast, disruption of the *trm11* gene caused growth retardation at high temperatures (35), and this temperature-sensitivity was further enhanced by the double disruption of *trm11* and *trm14* genes (Fig. 3H). Thus, Trm14- and Trm11-dependent modifications cooperate to maintain tRNA functions at high temperatures. To our knowledge, this represents the first example of a tRNA modification in the acceptor stem acting synergistically with the tertiary-core modifications m^2^G10/m^2^_2_G10 to promote cellular fitness.

Trm14 is composed of THUMP, NFLD, and Rossmann-fold methyltransferase domains. The THUMP domain is an RNA-binding domain found in a variety of RNA modification enzymes, including tRNA methyltransferases, thiouridine synthetases, deaminases, pseudouridine synthases, and a partner protein of tRNA acetyltransferase (12,14,15,18,31,36,47,55–59). The THUMP domain recognizes the CCA terminus of tRNA, which has been extensively characterized in previous studies (10,30,36,60–64). Furthermore, the NFLD is proposed to maintain the distance and orientation between the THUMP and catalytic domains, thereby determining the position of the target nucleotide within tRNA. Consequently, THUMP-family enzymes have been proposed to function as molecular rulers that recognize a specific nucleotide at a defined distance from the CCA terminus in tRNA (14,30,62–64). The finding that archaeal Trm14 methylates both G6 and G67 is therefore unexpected and raises the question of how a single THUMP-family enzyme recognizes two distinct target sites within tRNA. To address this question, we characterized the enzymatic properties of *T. kodakarensis* Trm14 in detail. Our analyses revealed that methylation efficiency is not determined by whether the target nucleotide is G6 or G67. Instead, local thermodynamic stability around the target site strongly influences Trm14 activity. Consistent with this interpretation, disruption of the target base pair accelerated the methylation reaction of Trm14, suggesting that Trm14 does not recognize the paired state of the target guanosine. Furthermore, dual-site specificity was found to be conserved among archaeal Trm14 enzymes but not in the bacterial ortholog TrmN.

Based on these findings and previous studies, we propose a potential mechanism for the archaeal Trm14 reaction (Fig. 9). First, we compared the geometric positions of G6 and G67 in the L-shaped tRNA structure (Fig. 9A and 9B). A previously proposed TrmN–tRNA complex model suggested that the enzyme accesses the target site from the minor groove side of the acceptor stem (17). We therefore examined the spatial arrangement of G6 and G67 using the tRNA^Phe^ structure (PDB ID: 1EHZ), in which U6 and A67 were replaced with G6 and G67, respectively (Fig. 9A). Remarkably, the distance between the *N*^2^ atoms of G6 (red ball in Fig. 9B) and G67 (blue ball in Fig. 9B) is only 1.3 Å, indicating that the two target atoms are located at nearly identical positions in three-dimensional space. Thus, binding of Trm14 to the minor groove of the acceptor stem may allow the enzyme to recognize the *N*^2^ atom of both G6 and G67, consistent with the TrmN-tRNA model (17). Alternatively, Trm14 may adopt multiple tRNA-binding modes that enable dual-site specificity. Furthermore, the TrmN–tRNA complex model strongly suggested that the target guanosine is flipped out from the acceptor stem and inserted into the catalytic pocket for methylation. Our biochemical analyses using mutant tRNA transcripts support this model (Figs. 5 and 6). Replacement of neighboring G-C base pairs with A-U base pairs, as well as disruption of the target G-C base pair itself, increased methylation efficiency. These substitutions are expected to enhance local breathing of the acceptor stem, thereby probably lowering the energetic barrier for capture and insertion of the target guanosine into the catalytic pocket. These observations support a flip-out mechanism for archaeal Trm14.

**Figure 9.**
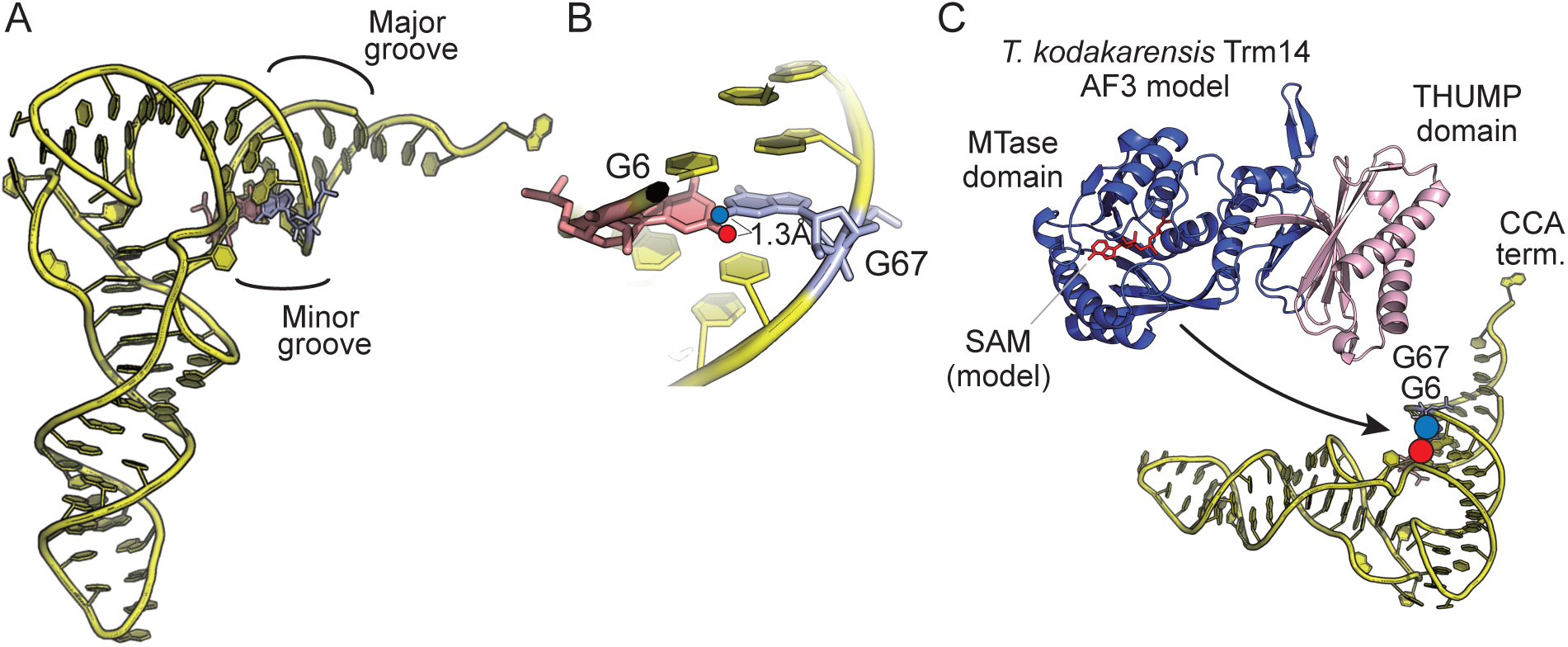
Structural model for dual-site recognition by archaeal Trm14. (A) Positions of G6 and G67 in the L-shaped tRNA structure are shown. The structure of yeast tRNA^Phe^ (PDB ID: 1EHZ) was used as a model, in which U6 and A67 were replaced with G6 and G67, respectively. The major and minor grooves of the acceptor stem are indicated. (B) Close-up view of the target nucleotides. The distance between the *N*^2^ atoms of G6 (red ball) and G67 (blue ball) is approximately 1.3 Å. (C) Proposed model of substrate recognition by *T. kodakarensis* Trm14.

Taken together, we summarized the Trm14 reaction (Fig. 9C). We modeled the *T. kodakarensis* Trm14 structure using AlphaFold3 (65). The THUMP domain is reported to recognize the CCA terminus of tRNA. The methyltransferase domain is proposed to bind to substrate tRNAs from the minor groove side to recognize both G6 and G67 (Fig. 9C).

In summary, our biochemical, genetic, and structural analyses characterized Trm14 as a dual-site specific tRNA m^2^G methyltransferase and demonstrated that m^2^G6/m^2^G67 and m^2^G10/m^2^_2_G10 cooperatively contribute to cellular fitness at high temperatures.

## Acknowledgements

The authors thank Dr. Kohei Nishino at Tokushima University for assistance with the nucleoside analyses. We are grateful to the lab member for helpful discussion.

## Author contributions

Teppei Matsuda (Investigation [lead], Formal Analysis [lead], Validation [lead], Visualization [lead], Funding acquisition [equal], Writing– review & editing [equal]), Takashi Yokogawa (Investigation [equal], Formal Analysis [equal], Validation [equal]), Soichiro Hidetaka (Investigation [supporting]), Manaka Sora (Investigation [supporting]), Aoi Ihara (Investigation [supporting]), Ayumu Toba (Investigation [supporting]), Kumpei Kawai (Investigation [supporting]), Go Norimoto (Investigation [supporting]), Akira Hirata (Conceptualization [equal], Funding acquisition [equal], Resources [lead], Investigation [equal], Supervision [equal], Formal Analysis [equal], Validation [equal], Visualization [equal], Writing–review & editing [equal]), Hiroyuki Hori (Conceptualization [lead], Funding acquisition [equal], Project administration [equal], Supervision [equal], Formal Analysis [equal], Validation [equal], Writing–review & editing [equal]), Ryota Yamagami (Conceptualization [equal], Funding acquisition [lead], Project administration [lead], Supervision [lead], Formal Analysis [equal], Validation [equal], Visualization [equal], Writing–original draft [lead], Writing–review & editing [lead]).

## Supplementary data

Supplementary data is available at BioRxiv Online.

## Conflict of interest

The authors declare no competing interests.

## Funding

This work was supported by the JAPAN Society for the Promotion of Science (JSPS) (24K09352 for A.H., 24K09381 for H.H., and 26KJ1761 for T.M.), funding from the Institute for Fermentation, Osaka, Grant Number LA-2026-006 to A.H. and G-2024-2- 059 to R.Y., the Uehara Memorial Foundation (202410178 for R.Y.), and the Sasakawa Scientific Research Grant from The Japan Science Society (2025-4046 for T.M.).

## Data Availability

All data used in this study are available from the corresponding author upon request.

## Notes

### Competing Interest Statement

The authors have declared no competing interest.

## References

1. Orellana, E.A., Siegal, E. and Gregory, R.I. (2022) tRNA dysregulation and disease. Nat. Rev. Genet., 23, 651–664.

2. Chujo, T. and Tomizawa, K. (2025) Neurological Diseases Caused by Loss of Transfer RNA Modifications: Commonalities in Their Molecular Pathogenesis. J Mol Biol, 437, 169047.

3. Stanley, R.E., Lowe, T.M. and Ignatova, Z. (2026) The regulation, function and disease relevance of cytoplasmic tRNAs. Nat Rev Mol Cell Biol.

4. Sordyl, D., Boileau, E., Bernat, A., Maiti, S., Mukherjee, S., Moafinejad, S.N., Farsani, M.A., Shavina, A., Cappannini, A., Agostini, G. et al. (2026) MODOMICS: a database of RNA modifications and related information. 2025 update and 20th anniversary. Nucleic Acids Res, 54, D219–D225.

5. McCloskey, J.A., Graham, D.E., Zhou, S., Crain, P.F., Ibba, M., Konisky, J., Soll, D. and Olsen, G.J. (2001) Post-transcriptional modification in archaeal tRNAs: identities and phylogenetic relations of nucleotides from mesophilic and hyperthermophilic Methanococcales. Nucleic Acids Res, 29, 4699–4706.

6. Hori, H., Kawamura, T., Awai, T., Ochi, A., Yamagami, R., Tomikawa, C. and Hirata, A. (2018) Transfer RNA modification enzymes from thermophiles and their modified nucleosides in tRNA. Microorganisms, 6, 110–152.

7. Lorenz, C., Lunse, C.E. and Mörl, M. (2017) tRNA Modifications: Impact on Structure and Thermal Adaptation. Biomolecules., 7, 35.

8. Suzuki, T. (2021) The expanding world of tRNA modifications and their disease relevance. Nat Rev Mol Cell Biol, 22, 375–392.

9. Ohira, T. and Suzuki, T. (2024) Transfer RNA modifications and cellular thermotolerance. Mol Cell, 84, 94–106.

10. Nishida, Y., Ohmori, S., Kakizono, R., Kawai, K., Namba, M., Okada, K., Yamagami, R., Hirata, A. and Hori, H. (2022) Required Elements in tRNA for Methylation by the Eukaryotic tRNA (Guanine-N(2)-) Methyltransferase (Trm11-Trm112 Complex). Int J Mol Sci, 23.

11. Leavitt, J.S., Moore, H.T., Santangelo, T.J. and Lowe, T.M. (2026) OTTR-seq profiling reveals dynamic tRNA modification landscapes across diverse archaeal species. Genome Biol, in press.

12. Menezes, S., Gaston, K.W., Krivos, K.L., Apolinario, E.E., Reich, N.O., Sowers, K.R., Limbach, P.A. and Perona, J.J. (2011) Formation of m^2^G6 in *Methanocaldococcus jannaschii* tRNA catalyzed by the novel methyltransferase Trm14. Nucleic Acids Res, 39, 7641–7655.

13. Aravind, L. and Koonin, E.V. (2001) THUMP--a predicted RNA-binding domain shared by 4- thiouridine, pseudouridine synthases and RNA methylases. Trends Biochem Sci, 26, 215–217.

14. Hori, H. (2023) Transfer RNA Modification Enzymes with a Thiouridine Synthetase, Methyltransferase and Pseudouridine Synthase (THUMP) Domain and the Nucleosides They Produce in tRNA. Genes (Basel), 14, 382.

15. Roovers, M., Oudjama, Y., Fislage, M., Bujnicki, J.M., Versées, W. and Droogmans, L. (2012) The open reading frame TTC1157 of Thermus thermophilus HB27 encodes the methyltransferase forming N²-methylguanosine at position 6 in tRNA. RNA, 18, 815–824.

16. Yamagami, R., Tomikawa, C., Shigi, N., Kazayama, A., Asai, S., Takuma, H., Hirata, A., Fourmy, D., Asahara, H., Watanabe, K. et al. (2016) Folate-/FAD-dependent tRNA methyltransferase from *Thermus thermophilus* regulates other modifications in tRNA at low temperatures. Genes Cells, 21, 740–754.

17. Fislage, M., Roovers, M., Tuszynska, I., Bujnicki, J.M., Droogmans, L. and Versees, W. (2012) Crystal structures of the tRNA:m2G6 methyltransferase Trm14/TrmN from two domains of life. Nucleic Acids Res, 40, 5149–5161.

18. Yang, W.Q., Xiong, Q.P., Ge, J.Y., Li, H., Zhu, W.Y., Nie, Y., Lin, X., Lv, D., Li, J., Lin, H. et al. (2021) THUMPD3-TRMT112 is a m^2^G methyltransferase working on a broad range of tRNA substrates. Nucleic Acids Res, 49, 11900–11919.

19. Studte, P., Zink, S., Jablonowski, D., Bar, C., von der Haar, T., Tuite, M.F. and Schaffrath, R. (2008) tRNA and protein methylase complexes mediate zymocin toxicity in yeast. Mol Microbiol, 69, 1266–1277.

20. Figaro, S., Wacheul, L., Schillewaert, S., Graille, M., Huvelle, E., Mongeard, R., Zorbas, C., Lafontaine, D.L. and Heurgue-Hamard, V. (2012) Trm112 is required for Bud23-mediated methylation of the 18S rRNA at position G1575. Mol Cell Biol, 32, 2254–2267.

21. Letoquart, J., van Tran, N., Caroline, V., Aleksandrov, A., Lazar, N., van Tilbeurgh, H., Liger, D. and Graille, M. (2015) Insights into molecular plasticity in protein complexes from Trm9- Trm112 tRNA modifying enzyme crystal structure. Nucleic Acids Res, 43, 10989–11002.

22. Bourgeois, G., Letoquart, J., van Tran, N. and Graille, M. (2017) Trm112, a Protein Activator of Methyltransferases Modifying Actors of the Eukaryotic Translational Apparatus. Biomolecules, 7.

23. Wang, C., Ulryck, N., Herzel, L., Pythoud, N., Kleiber, N., Guerineau, V., Jactel, V., Moritz, C., Bohnsack, M.T., Carapito, C. et al. (2023) *N*^2^-methylguanosine modifications on human tRNAs and snRNA U6 are important for cell proliferation, protein translation and pre-mRNA splicing. Nucleic Acids Res, 51, 7496–7519.

24. Constantinesco, F., Benachenhou, N., Motorin, Y. and Grosjean, H. (1998) The tRNA(guanine- 26,*N*^2^-*N*^2^) methyltransferase (Trm1) from the hyperthermophilic archaeon P*yrococcus furiosus*: cloning, sequencing of the gene and its expression in *Escherichia coli*. Nucleic Acids Res, 26, 3753–3761.

25. Armengaud, J., Urbonavicius, J., Fernandez, B., Chaussinand, G., Bujnicki, J.M. and Grosjean, H. (2004) *N*^2^-methylation of guanosine at position 10 in tRNA is catalyzed by a THUMP domain-containing, S-adenosylmethionine-dependent methyltransferase, conserved in Archaea and Eukaryota. J Biol Chem, 279, 37142–37152.

26. Gabant, G., Auxilien, S., Tuszynska, I., Locard, M., Gajda, M.J., Chaussinand, G., Fernandez, B., Dedieu, A., Grosjean, H., Golinelli-Pimpaneau, B. et al. (2006) THUMP from archaeal tRNA:m^2^_2_G10 methyltransferase, a genuine autonomously folding domain. Nucleic Acids Res, 34, 2483–2494.

27. Grosjean, H., Gaspin, C., Marck, C., Decatur, W.A. and de Crecy-Lagard, V. (2008) RNomics and Modomics in the halophilic archaea Haloferax volcanii: identification of RNA modification genes. BMC Genomics, 9, 470.

28. Ihsanawati, Nishimoto, M., Higashijima, K., Shirouzu, M., Grosjean, H., Bessho, Y. and Yokoyama, S. (2008) Crystal structure of tRNA *N*^2^,*N*^2^-guanosine dimethyltransferase Trm1 from *Pyrococcus horikoshii*. J Mol Biol, 383, 871–884.

29. Awai, T., Kimura, S., Tomikawa, C., Ochi, A., Ihsanawati, Bessho, Y., Yokoyama, S., Ohno, S., Nishikawa, K., Yokogawa, T., et al. (2009) Aquifex aeolicus tRNA (*N^2^*,*N^2^*-guanine)- dimethyltransferase (Trm1) catalyzes transfer of methyl groups not only to guanine 26 but also to guanine 27 in tRNA. J Biol Chem, 284, 20467–20478.

30. Hirata, A., Nishiyama, S., Tamura, T., Yamauchi, A. and Hori, H. (2016) Structural and functional analyses of the archaeal tRNA m^2^G/m^2^_2_G10 methyltransferase aTrm11 provide mechanistic insights into site specificity of a tRNA methyltransferase that contains common RNA-binding modules. Nucleic Acids Research, 44, 6377–6390.

31. Wang, C., van Tran, N., Jactel, V., Guerineau, V. and Graille, M. (2020) Structural and functional insights into Archaeoglobus fulgidus m^2^G10 tRNA methyltransferase Trm11 and its Trm112 activator. Nucleic Acids Res, 48, 11068–11082.

32. Shi, H. and Moore, P.B. (2000) The crystal structure of yeast phenylalanine tRNA at 1.93 A resolution: a classic structure revisited. RNA, 6, 1091–1105.

33. Yanagihara, K., Konishi, F., Matsuda, T., Hirata, A., Hori, H., Bevilacqua, P.C. and Yamagami, R. (2026) Optimized tRNA structure-seq reveals robust tRNA secondary structures in *S. cerevisiae* under mild stress conditions. RNA, in press.

34. Morikawa, M., Izawa, Y., Rashid, N., Hoaki, T. and Imanaka, T. (1994) Purification and characterization of a thermostable thiol protease from a newly isolated hyperthermophilic Pyrococcus sp. Appl Environ Microbiol, 60, 4559–4566.

35. Hirata, A., Suzuki, T., Nagano, T., Fujii, D., Okamoto, M., Sora, M., Lowe, T.M., Kanai, T., Atomi, H., Suzuki, T. et al. (2019) Distinct modified nucleosides in tRNA^Trp^ from the hyperthermophilic archaeon *Thermococcus kodakarensis* and requirement of tRNA m^2^G10/m^2^_2_G10 methyltransferase (archaeal Trm11) for survival at high temperatures. J Bacteriol, 201.

36. Matsuda, T., Yamagami, R., Ihara, A., Suzuki, T., Hirata, A. and Hori, H. (2025) A transfer RNA methyltransferase with an unusual domain composition catalyzes 2’-O-methylation at position 6 in tRNA. Nucleic Acids Res, 53, 1–19.

37. Sato, T., Fukui, T., Atomi, H. and Imanaka, T. (2003) Targeted gene disruption by homologous recombination in the hyperthermophilic archaeon *Thermococcus kodakaraensis* KOD1. J Bacteriol, 185, 210–220.

38. Atomi, H., Fukui, T., Kanai, T., Morikawa, M. and Imanaka, T. (2004) Description of *Thermococcus kodakaraensis* sp. nov., a well studied hyperthermophilic archaeon previously reported as *Pyrococcus* sp. KOD1. Archaea, 1, 263–267.

39. Sato, T., Fukui, T., Atomi, H. and Imanaka, T. (2005) Improved and versatile transformation system allowing multiple genetic manipulations of the hyperthermophilic archaeon Thermococcus kodakaraensis. Appl Environ Microbiol, 71, 3889–3899.

40. Matsuda, T., Hori, H. and Yamagami, R. (2024) Rational design of oligonucleotides for enhanced in vitro transcription of small RNA. RNA, 30, 710–727.

41. Yokogawa, T., Kitamura, Y., Nakamura, D., Ohno, S. and Nishikawa, K. (2010) Optimization of the hybridization-based method for purification of thermostable tRNAs in the presence of tetraalkylammonium salts. Nucleic Acids Res, 38, e89.

42. Kazayama, A., Yamagami, R., Yokogawa, T. and Hori, H. (2015) Improved solid-phase DNA probe method for tRNA purification: large-scale preparation and alteration of DNA fixation. J. Biochem., 157, 411–418.

43. Chan, P.P. and Lowe, T.M. (2016) GtRNAdb 2.0: an expanded database of transfer RNA genes identified in complete and draft genomes. Nucleic Acids Res, 44, D184–189.

44. Xia, T., SantaLucia, J., Jr., Burkard, M.E., Kierzek, R., Schroeder, S.J., Jiao, X., Cox, C. and Turner, D.H. (1998) Thermodynamic parameters for an expanded nearest-neighbor model for formation of RNA duplexes with Watson-Crick base pairs. Biochemistry, 37, 14719–14735.

45. Mathews, D.H., Disney, M.D., Childs, J.L., Schroeder, S.J., Zuker, M. and Turner, D.H. (2004) Incorporating chemical modification constraints into a dynamic programming algorithm for prediction of RNA secondary structure. Proc Natl Acad Sci U S A, 101, 7287–7292.

46. Zuber, J., Schroeder, S.J., Sun, H., Turner, D.H. and Mathews, D.H. (2022) Nearest neighbor rules for RNA helix folding thermodynamics: improved end effects. Nucleic Acids Res, 50, 5251–5262.

47. Purushothaman, S.K., Bujnicki, J.M., Grosjean, H. and Lapeyre, B. (2005) Trm11p and Trm112p are both required for the formation of 2-methylguanosine at position 10 in yeast tRNA. Mol Cell Biol, 25, 4359–4370.

48. Alexandrov, A., Chernyakov, I., Gu, W., Hiley, S.L., Hughes, T.R., Grayhack, E.J. and Phizicky, E.M. (2006) Rapid tRNA decay can result from lack of nonessential modifications. Mol Cell, 21, 87–96.

49. Phizicky, E.M. and Hopper, A.K. (2010) tRNA biology charges to the front. Genes Dev, 24, 1832–1860.

50. Okamoto, M., Fujiwara, M., Hori, M., Okada, K., Yazama, F., Konishi, H., Xiao, Y., Qi, G., Shimamoto, F., Ota, T. et al. (2014) tRNA modifying enzymes, NSUN2 and METTL1, determine sensitivity to 5-fluorouracil in HeLa cells. PLoS Genet, 10, e1004639.

51. Kotelawala, L., Grayhack, E.J. and Phizicky, E.M. (2008) Identification of yeast tRNA Um(44) 2’-*O*-methyltransferase (Trm44) and demonstration of a Trm44 role in sustaining levels of specific tRNA^Ser^ species. RNA, 14, 158–169.

52. Kimura, S. and Waldor, M.K. (2019) The RNA degradosome promotes tRNA quality control through clearance of hypomodified tRNA. Proceedings of the National Academy of Sciences, 116, 1394–1403.

53. Schultz, S.K. and Kothe, U. (2024) RNA modifying enzymes shape tRNA biogenesis and function. J Biol Chem, 300, 107488.

54. Schultz, S.K., Katanski, C.D., Halucha, M., Pena, N., Fahlman, R.P., Pan, T. and Kothe, U. (2024) Modifications in the T arm of tRNA globally determine tRNA maturation, function, and cellular fitness. Proc Natl Acad Sci U S A, 121, e2401154121.

55. Johansson, M.J. and Bystrom, A.S. (2004) The *Saccharomyces cerevisiae* TAN1 gene is required for *N*^4^-acetylcytidine formation in tRNA. RNA, 10, 712–719.

56. Waterman, D.G., Ortiz-Lombardia, M., Fogg, M.J., Koonin, E.V. and Antson, A.A. (2006) Crystal structure of Bacillus anthracis ThiI, a tRNA-modifying enzyme containing the predicted RNA-binding THUMP domain. J Mol Biol, 356, 97–110.

57. Roovers, M., Hale, C., Tricot, C., Terns, M.P., Terns, R.M., Grosjean, H. and Droogmans, L. (2006) Formation of the conserved pseudouridine at position 55 in archaeal tRNA. Nucleic Acids Res, 34, 4293–4301.

58. Randau, L., Stanley, B.J., Kohlway, A., Mechta, S., Xiong, Y. and Soll, D. (2009) A cytidine deaminase edits C to U in transfer RNAs in Archaea. Science, 324, 657–659.

59. Sharma, S., Langhendries, J.L., Watzinger, P., Kotter, P., Entian, K.D. and Lafontaine, D.L. (2015) Yeast Kre33 and human NAT10 are conserved 18S rRNA cytosine acetyltransferases that modify tRNAs assisted by the adaptor Tan1/THUMPD1. Nucleic Acids Res, 43, 2242–2258.

60. Lauhon, C.T., Erwin, W.M. and Ton, G.N. (2004) Substrate specificity for 4-thiouridine modification in Escherichia coli. J Biol Chem, 279, 23022–23029.

61. Tanaka, Y., Yamagata, S., Kitago, Y., Yamada, Y., Chimnaronk, S., Yao, M. and Tanaka, I. (2009) Deduced RNA binding mechanism of ThiI based on structural and binding analyses of a minimal RNA ligand. RNA, 15, 1498–1506.

62. Neumann, P., Lakomek, K., Naumann, P.T., Erwin, W.M., Lauhon, C.T. and Ficner, R. (2014) Crystal structure of a 4-thiouridine synthetase-RNA complex reveals specificity of tRNA U8 modification. Nucleic Acids Res, 42, 6673–6685.

63. Bourgeois, G., Marcoux, J., Saliou, J.M., Cianferani, S. and Graille, M. (2017) Activation mode of the eukaryotic m^2^G10 tRNA methyltransferase Trm11 by its partner protein Trm112. Nucleic Acids Research, 45, 1971–1982.

64. Ma, C.R., Liu, N., Li, H., Xu, H. and Zhou, X.L. (2024) Activity reconstitution of Kre33 and Tan1 reveals a molecular ruler mechanism in eukaryotic tRNA acetylation. Nucleic Acids Res, 52, 5226–5240.

65. Abramson, J., Adler, J., Dunger, J., Evans, R., Green, T., Pritzel, A., Ronneberger, O., Willmore, L., Ballard, A.J., Bambrick, J. et al. (2024) Accurate structure prediction of biomolecular interactions with AlphaFold 3. Nature, 630, 493–500.

