## Supplementary Figures for "Dual-site specificity of the archaeal tRNA m^2^G methyltransferase Trm14"

**Running title: Dual-site specificity of archaeal Trm14**

**Keywords:** Archaea, tRNA, tRNA methyltransferase, tRNA methylation, dual-site specificity

**Supplementary Figure 1**

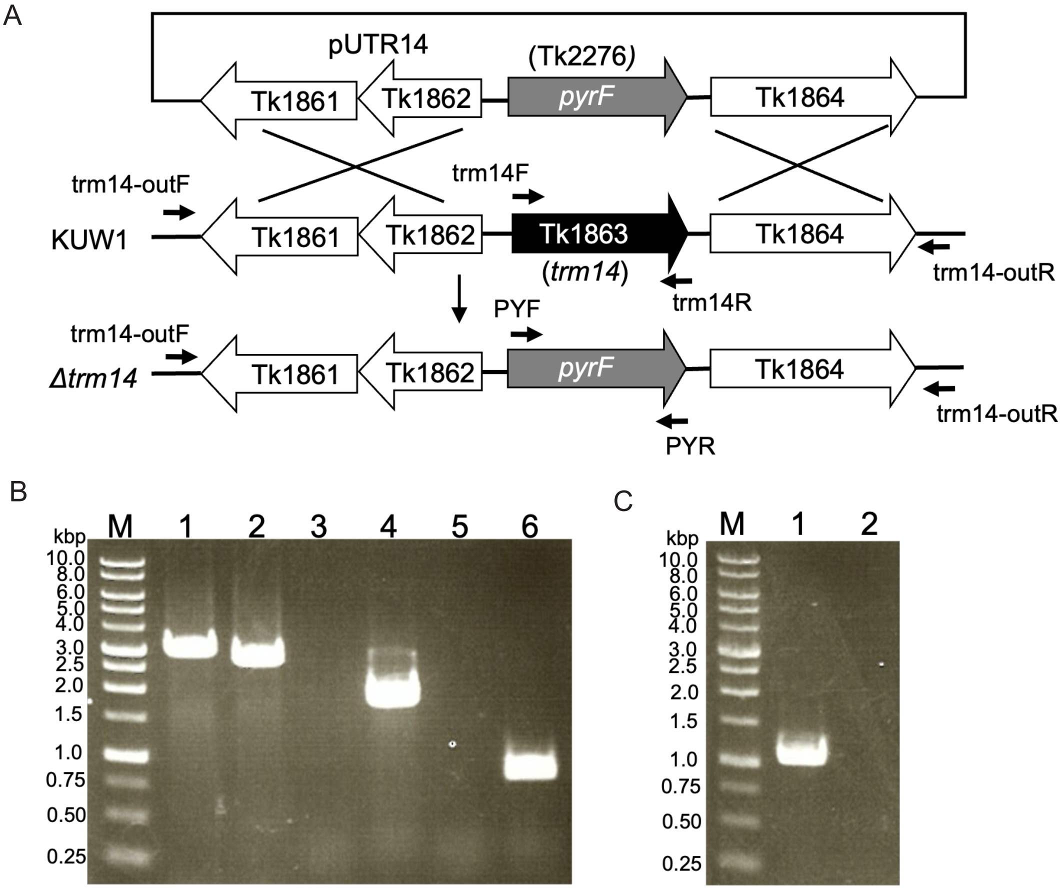


**Construction of the *Δtrm14* strain.** (A) The KUW1 strain was used as the parent strain and homologous recombination with a plasmid vector, pUTR14, in which the selectable marker gene (*pyrF*; Tk2276) was inserted between the Tk1862 and Tk1864. The vector, pUTR14, is not maintained as a plasmid in *T. kodakarensis* cells due to the lack of a replication origin. The pairs of oligonucleotides (trm14-outF and PYR; sequences in Table S2) used to amplify genomic DNA from pUTR14-generated uracil-independent transformants are indicated. (B) Genomic DNA from *T. kodakarensis* KUW1 (lanes 1, 3, and 5) and Δ*trm14* (lanes 2, 4, and 6) was amplified by PCR using different primer combinations (primer sequences are listed in Table S2). Lane 1: KUW1 DNA amplified with primers trm14-outF and trm14-outR, yielding a ~3.0 kb fragment. Lane 2: *Δtrm14* DNA amplified with trm14-outF and trm14-outR, yielding a ~2.7 kb fragment. Lane 3: KUW1 DNA amplified with trm14-outF and PYR, no detectable PCR product. Lane 4: *Δtrm14* DNA amplified with trm14-outF and PYR, yielding a 1.7 kb fragment. Lane 5: KUW1 DNA amplified with PYF and PYR, no detectable PCR product. Lane 6: *Δtrm14* DNA amplified with PYF and PYR, yielding a 0.75 kb fragment. M, DNA size marker. (C) PCR amplification using primers trm14F and trm14R. Lane 1: KUW1 genomic

DNA, yielding a ~1.0 kb fragment. Lane 2: *Δtrm14* genomic DNA, showing no detectable amplification product. M, DNA size marker.

**Supplementary Figure 2**

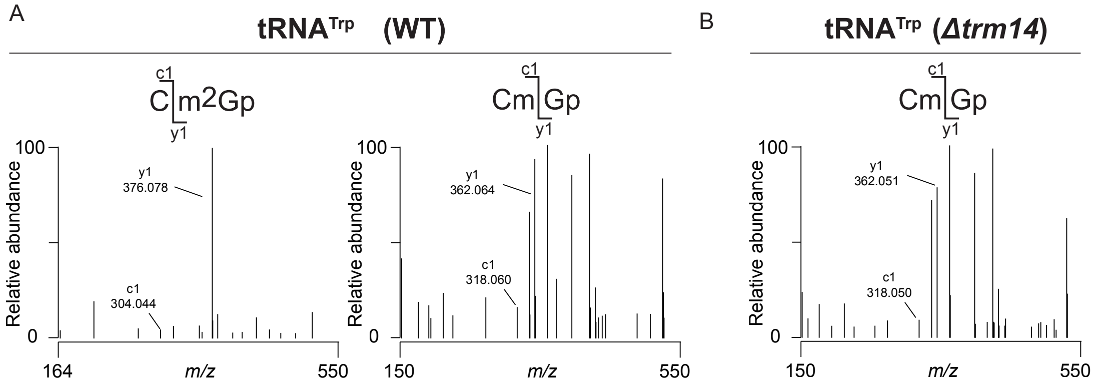


**LC–MS/MS analyses of RNase T1-digested fragments of native tRNA$^{Trp}$**

RNase T1-digested fragments of tRNA$^{Trp}$ from WT and Δ*trm14* strains were analyzed by LC–MS/MS. Extracted ion chromatograms of RNase T1-digested fragments (m/z 681.115, z = -1) are shown in Fig. 3C. Detected peaks were further analyzed by MS/MS. (A) CID spectra of 5'-Cm$^2$Gp-3' and 5'-CmGp-3' fragments of native tRNA$^{Trp}$ purified from the WT (KUW1) strain are shown. (B) CID spectrum of the 5'-CmGp-3' fragment of native tRNA$^{Trp}$ purified from the Δ*trm14* strain is shown.

**Supplementary Figure 3**

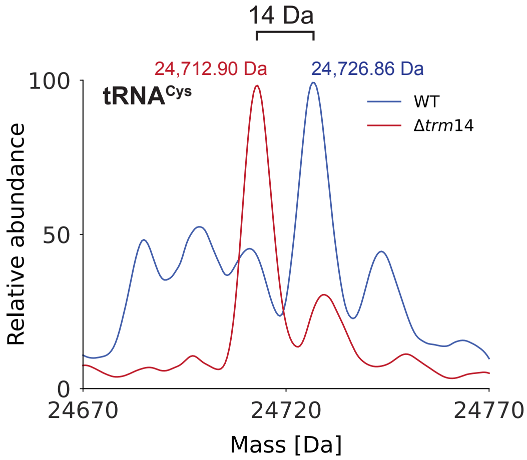


**LC–MS analysis of intact tRNA$^{Cys}$ purified from the WT and Δ*trm14* strains**

Molecular weights of the intact tRNA$^{Cys}$ purified from the WT and Δ*trm14* strains were measured by LC–MS. The tRNA from the WT and Δ*trm14* strains are highlighted in blue and red, respectively.

**Supplementary Figure 4**

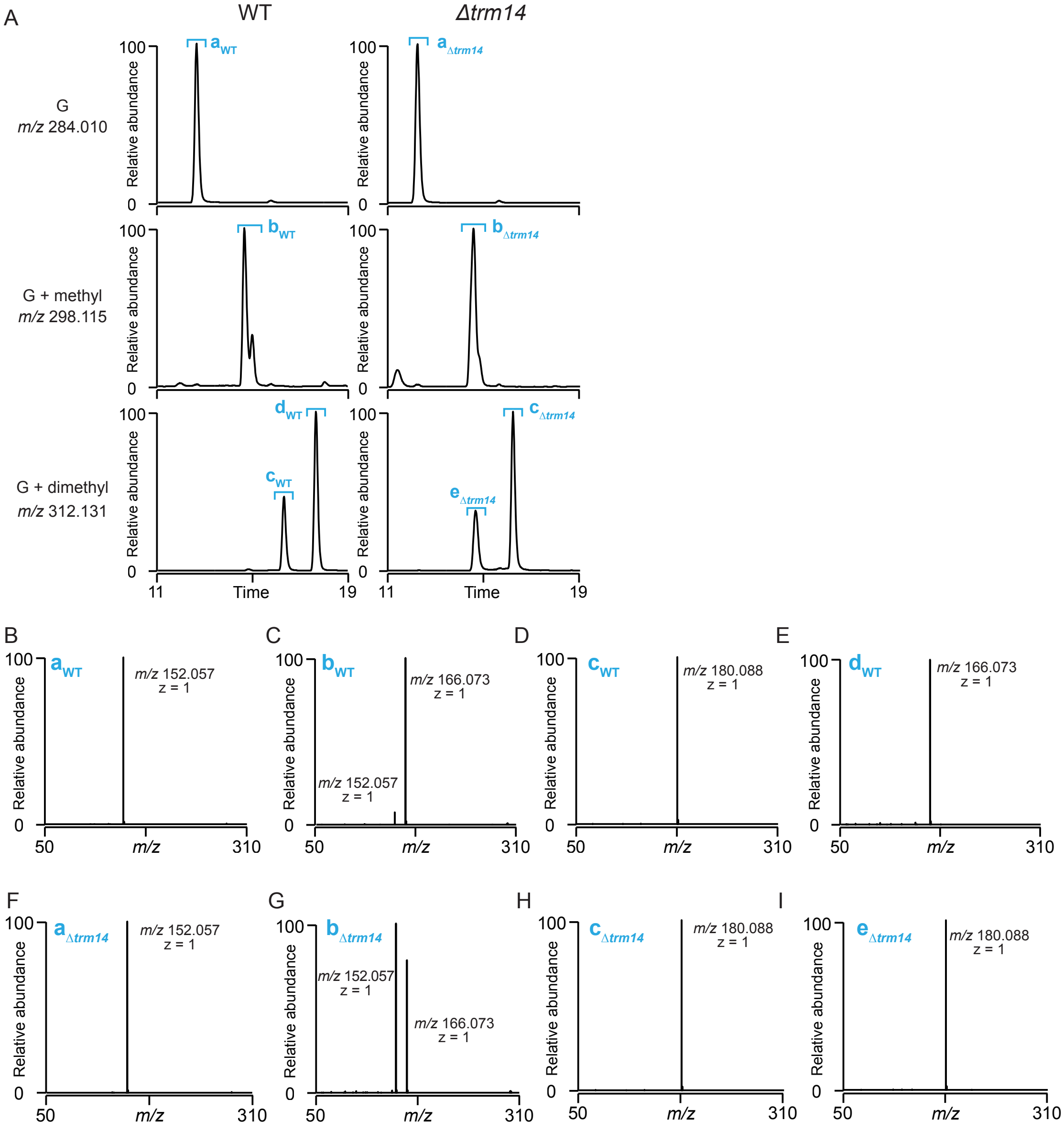


**Nucleoside analyses of native tRNA$^{Cys}$ from the WT and Δ*trm14* strains.**

(A) Extracted ion chromatograms of guanosine (m/z 284.010, z = +1), mono-methylated guanosine (m/z 298.115, z = +1), and di-methylated guanosine (m/z 312.131, z = +1) are shown. Each peak was labeled alphabetically. (B-I) The peaks detected in the extracted ion chromatograms were further analyzed by MS/MS. The m/z values and charge states of the product ions are shown. (B and F) CID spectra of the peaks assigned as $a_{WT}$ and $a_{\Delta trm14}$ in panel (A). (C and G) CID spectra of the peaks assigned as $b_{WT}$ and $b_{\Delta trm14}$ in panel (A). (D and H) CID spectra of the peaks assigned as $c_{WT}$ and $c_{\Delta trm14}$ in panel (A). (E and I) CID spectra of the peaks assigned as $d_{WT}$ and $e_{\Delta trm14}$ in panel (A). The peak $e_{\Delta trm14}$ corresponds to the peak labeled with the asterisk in main Figure 3F.

**Supplementary Figure 5**

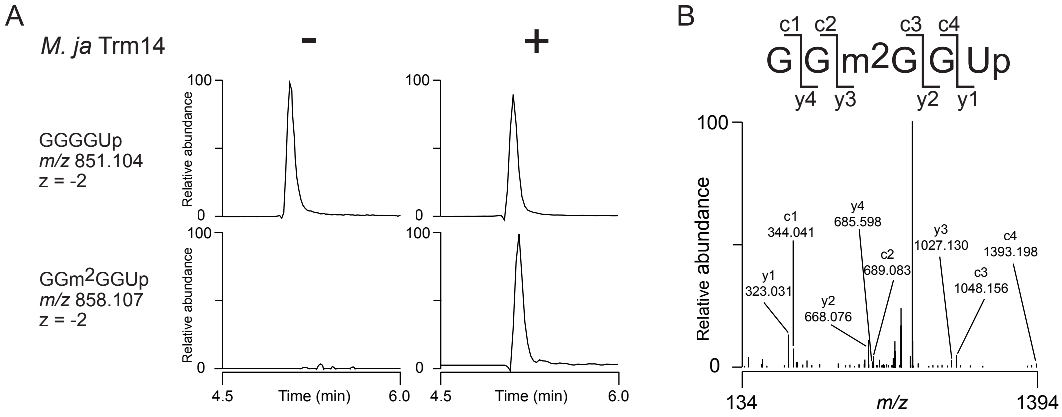


**LC–MS/MS analysis of *M. jannaschii* tRNA$^{Cys}$ transcript**

(A) Extracted ion chromatograms of the indicated negative ions derived from RNase A-digested tRNA$^{Cys}$ transcripts reacted with or without recombinant *M. jannaschii* Trm14 are shown. (B) CID spectrum of the m$^2$G-containing fragment from the Trm14-methylated tRNA$^{Cys}$ transcript.

### Supplementary Figure 6

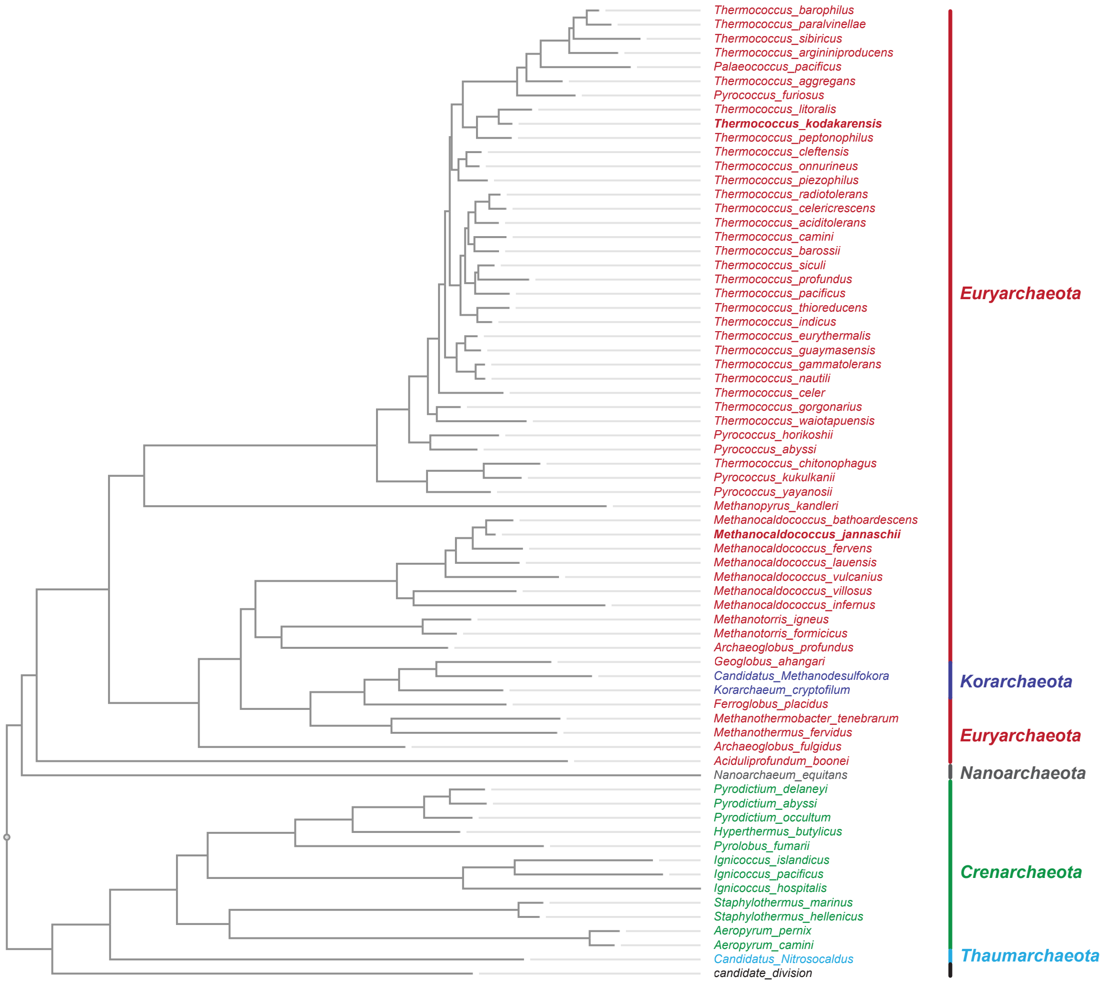


**Phylogenetic analysis of Trm14-like proteins.** Trm14-like proteins were identified by homology searching using *T. kodakarensis* Trm14 (TK1863) as the query sequence. Proteins with BLAST E-values < 1 x $10^{-30}$ were retrieved and aligned using MAFFT with default parameters. A phylogenetic tree was constructed using the neighbor-joining (NJ) method from the resulting multiple sequence alignment. The tree was visualized using Archaeopteryx.js. Species names are colored according to their archaeal phyla: Euryarchaeota (red), Korarchaeota (blue), Nanoarchaeota (gray), Crenarchaeota (green), Thaumarchaeota (light blue), and archaeal candidate divisions (black). *T. kodakarensis* and *M. jannaschii* were bolded.

**Supplementary Figure 7**

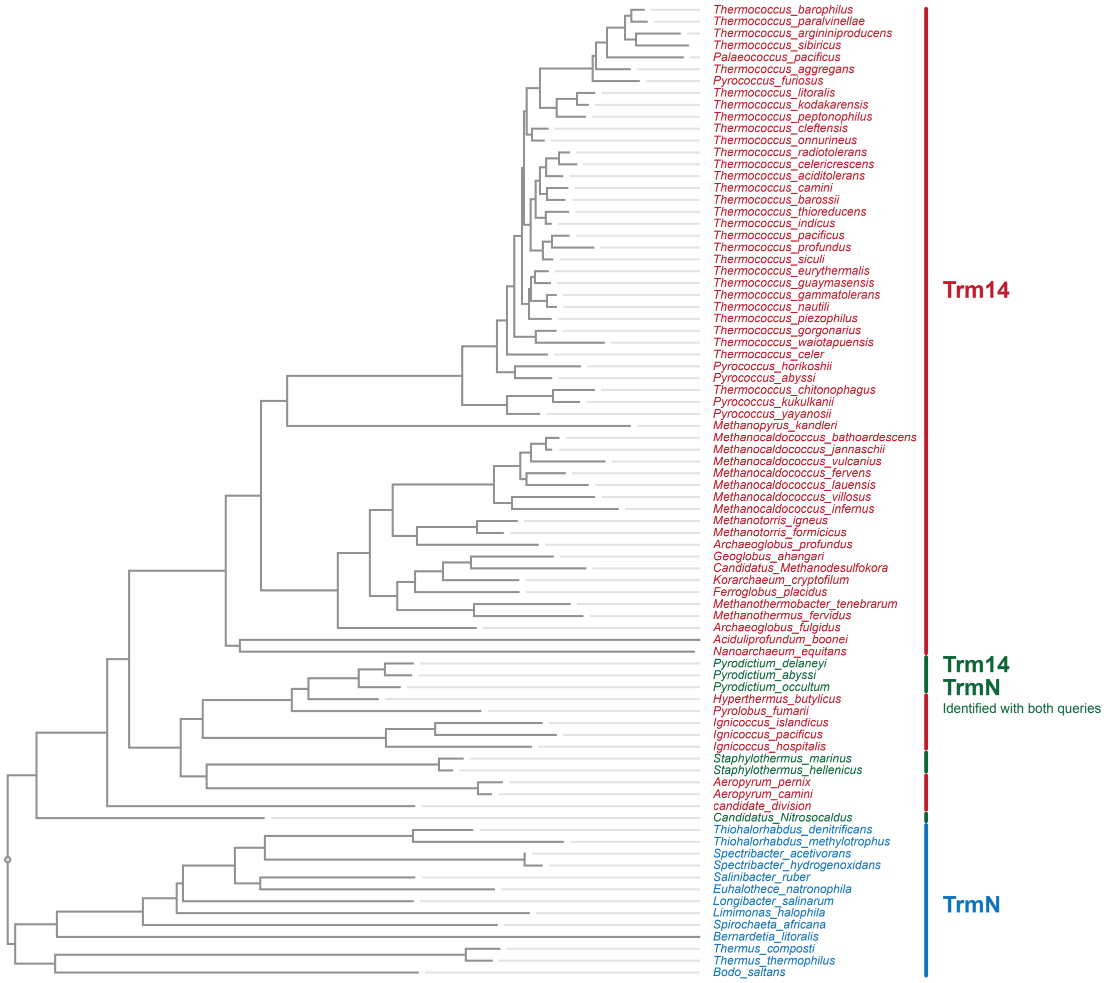


**Phylogenetic analysis of Trm14- and TrmN-like proteins.** Trm14- and TrmN-like proteins were independently identified by homology searches using *T. kodakarensis* Trm14 (TK1863) or *T. thermophilus* HB27 TrmN (TTC1157) as the query sequence. Proteins with BLAST E-values < $1 \times 10^{-30}$ were retrieved, pooled, and aligned using MAFFT with default parameters. A phylogenetic tree was constructed using the neighbor-joining (NJ) method from the resulting multiple sequence alignment. Organism names are colored according to the homology search by which they were identified: red, proteins identified only by the Trm14 query; blue, proteins identified only by the TrmN query; and green, proteins identified by both Trm14 and TrmN queries.
